# Ferritinophagy is a Druggable Vulnerability of Quiescent Leukemic Stem Cells

**DOI:** 10.1101/2023.12.18.572101

**Authors:** Clement Larrue, Sarah Mouche, Paolo Angelino, Maxime Sajot, Rudy Birsen, François Vergez, Christian Recher, Véronique Mansat-De Mas, Qiong Gu, Jun Xu, Petros Tsantoulis, Jean-Emmanuel Sarry, Jerome Tamburini

## Abstract

Acute myeloid leukemia (AML) remains a challenging hematological malignancy with poor prognosis and limited treatment options. Leukemic stem cells (LSCs) contributes to therapeutic failure, post-therapy relapse and adverse outcome. Here, we investigated the role of quiescence and its associated molecular mechanisms in AML pathogenesis and LSCs functions, and identified potential vulnerabilities for therapeutic intervention. We found that LSC-enriched quiescent cell population exhibited a distinct gene set of prognostic significance in AML patients. Furthermore, this quiescent cells subset displayed heightened autophagic activity with a reliance on ferritinophagy, a selective form of autophagy mediated by Nuclear Receptor Coactivator 4 (NCOA4) regulating iron bioavailability. Inhibition of NCOA4 genetically or chemically showed potent anti-leukemic effects, particularly targeting the LSC compartment. These findings uncover that ferritinophagy inhibition may represent a promising therapeutic strategy for patients with AML.

**One Sentence Summary:** Targeting quiescent leukemic stem cells via NCOA4-dependent ferritinophagy inhibition may improve therapeutic outcomes in acute myeloid leukemia.

## INTRODUCTION

Acute myeloid leukemia (AML) is a hematological malignancy characterized by the rapid proliferation of immature myeloid cells. Despite recent therapeutic advances, AML remains associated with a poor prognosis and high mortality rate (*1*). Chemotherapy is the mainstay of treatment for patients with AML, but its efficacy is often limited by the emergence of drug-resistant cells leading to disease relapse, progression, and poor outcomes (*2*). One of the major contributors to chemoresistance is the presence of leukemic stem cells (LSCs), which are responsible for the initiation, propagation, and maintenance of AML (*3*).

Recent studies have shown that LSCs possess the capacity to self-renew and differentiate into various leukemic cells, and are characterized by their ability to survive chemotherapy and regenerate the disease upon relapse (*4–6*). One mechanism that allows LSCs to survive and evade therapy is their ability to enter and exit a quiescent state, in which they temporarily cease cell division and become refractory to cytotoxic treatments (*5*). This mechanism is shared between LSCs and normal hematopoietic stem cells (HSCs) and involves several processes such as the expression of cell cycle inhibitors, activation of signaling pathways, epigenetic modifications, and metabolic changes (*7*).

Iron is an essential nutrient that is indispensable for fundamental cellular functions, including DNA synthesis, mitochondrial metabolism and cell proliferation, and plays a critical role in the survival and expansion of cancer cells (*8*). Iron is predominantly obtained from dietary intake. The hormone hepcidin inhibit the activity of the iron transporter ferroportin, thereby reducing iron absorption. Iron is transported in the blood bound to transferrin, which delivers iron to cells expressing transferrin receptors. When not employed for cellular functions, excess iron is stored in ferritin to prevent toxicity (*9*). Nuclear Receptor Coactivator 4 (NCOA4) is a crucial effector of ferritinophagy, a selective form of autophagy that targets ferritin to the autophagosome for degradation, allowing the release of iron that can be used for cellular processes (*10*). Interestingly, recent studies found that inhibition of NCOA4 results in delayed tumor growth and prolonged survival in murine models of pancreatic cancer, while enhanced ferritinophagy accelerates tumorigenesis (*11, 12*).

Here, we investigated quiescent cell subpopulations relative to their cycling counterpart in patient-derived xenograft (PDX) models (*13*) and primary tumors from patients with AML. Our results revealed that quiescent AML cells displayed enrichment of LSC functions and carried a distinct gene set with significant prognostic implications for poor outcomes in AML patients. Furthermore, autophagic activity was heightened in quiescent cells, playing a crucial role in maintaining in iron bioavailability in this subset. Remarkably, quiescent cells relied on ferritinophagy, and blocking NCOA4 genetically or chemically exhibited anti-leukemic effects, particularly targeting the LSC compartment. Our results implicate ferritinophagy as a novel vulnerability of quiescent LSCs, underscoring its potential as a promising therapeutic target for patients with AML.

## RESULTS

### In Vivo Labeling Assays Reveal Differential Trajectories of Quiescent and Cycling Cells in AML PDX Models

Pioneering investigations on cell kinetics performed in patients with AML uncovered that only a minority of leukemic cells were actively dividing within the bone marrow, with an even smaller proportion in the bloodstream (*14, 15*). In line with these findings, previous investigations carried out in patient-derived xenograft (PDX) models have unveiled that a substantial subset of human leukemia cells reside in a quiescent state (*16–18*). Here, we aimed to expand these findings and delineate the evolution of quiescent and cycling cell subpopulations through in vivo labelling assays. To achieve this objective, we xenografted five different primary AML specimens from patients (**Table 1**) into immunodeficient NOD/SCID/IL-2R-γchain null (NSG) mice through two consecutive rounds of serial transplantation. This allowed us to establish a substantial reserve of leukemia cells that could be utilized for diverse ex vivo and in vivo assays (**Figure 1A**). We differentially probed human and murine hematopoietic cells with CD45 staining on mice bone marrow samples, and confirmed that most cells were viable human CD45^+^CD33^+^ leukemic cells (**Figure S1A-B**). We subsequently monitored the cell cycle distribution of these AML cells, and observed that 48.9%, 39.2% and 11.4% of cells were in G_0_ (Ki67^-^DAPI^-^), G_1_ (Ki67^+^DAPI^-^) and S/G_2_/M (Ki67^+^DAPI^+^) phases, respectively (**Figure 1B-C** and **Figure S1C**). Notably, the proportion of cells in the S/G_2_/M phase was comparable to that observed in bone marrow samples from AML patients at diagnosis (*15*).

**Figure 1.**
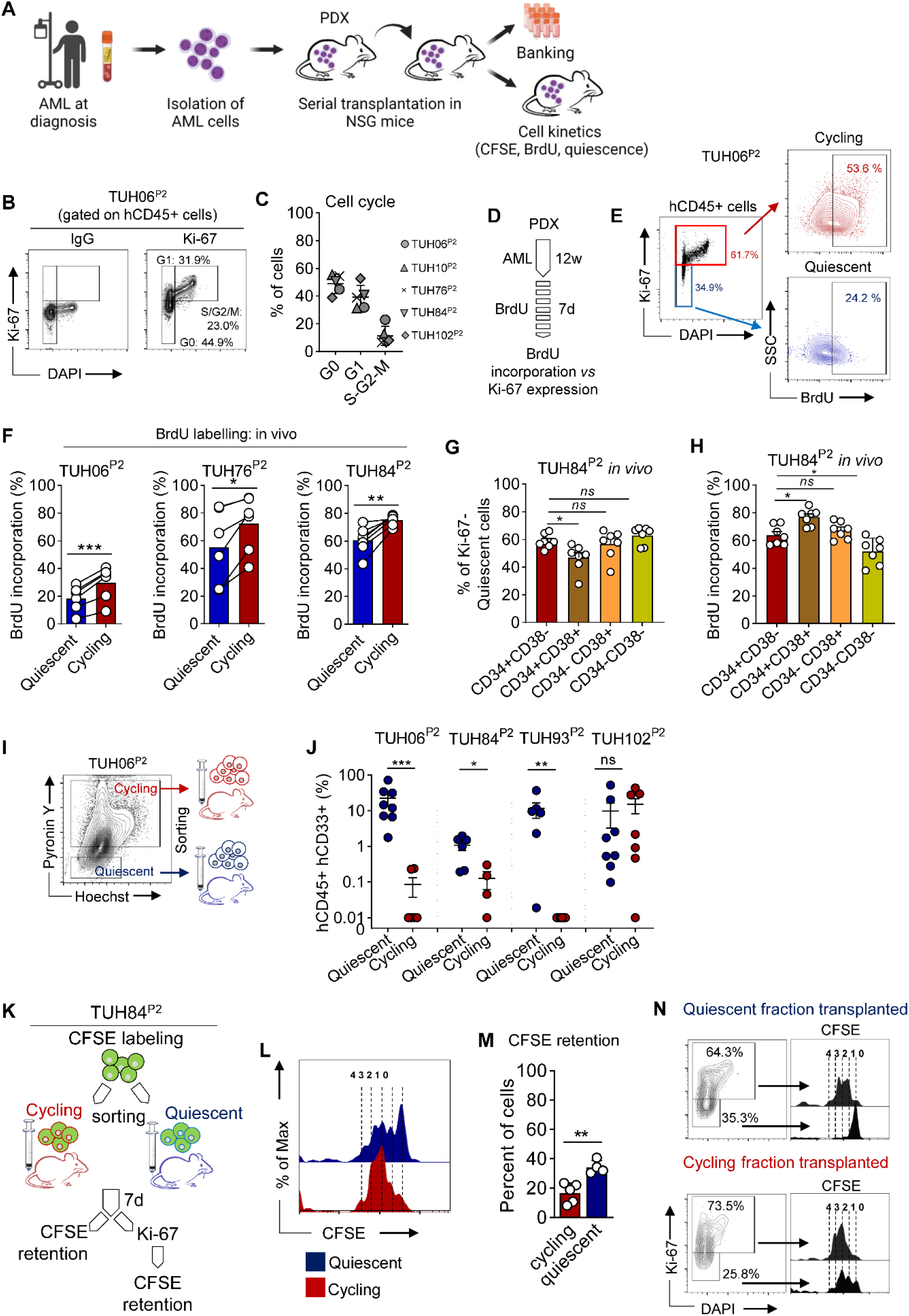
In Vivo Labeling Assays Reveal Differential Trajectories of Quiescent and Cycling Cells in AML PDX Models. **A.** Serial transplantation of primary AML specimens into immunodeficient NSG mice established a substantial reservoir of leukemia cells, enabling a wide range of assays, including in vivo cell kinetic experiments. **B.** Cell cycle analysis of human CD45^+^ leukemic cells was conducted using DAPI and staining with IgG (isotype) or Ki67. **C.** Distribution of cells within the G_0_, G_1_, and S-G_2_-M phases was determined in patient-derived xenograft (PDX) assays (n=5). **D.** Mice harboring AML PDXs were treated with 100mg/kg BrdU twice a day via intraperitoneal injections for seven consecutive days. BrdU incorporation was evaluated by staining CD45^+^ human cells isolated from the mice’s bone marrow with an anti-BrdU antibody. Quantification of BrdU incorporation was performed in distinct subpopulations based on their cell cycle status, including quiescent (Ki67^-^) and cycling cells (Ki67^+^). **E.** Representative contour plots illustrate BrdU labeling in quiescent or cycling human hCD45^+^ AML cells. **F.** The percentage of BrdU incorporation in quiescent versus cycling subpopulations was determined in vivo (n=6-7 mice per group in three different PDX assays). **G-H**. Human leukemic cells were stained with CD34 and CD38 antibodies. The proportion of quiescent (**G**) and BrdU^+^ (**H**) cells was examined within each subpopulation defined by CD34/CD38 staining (n=7 mice per group in one PDX assay). **I-J.** Cycling or quiescent populations were sorted using pyronin Y and Hoechst and subsequently transplanted separately into NSG mice. The schematic representation of the experimental setup is shown (**I**). **J.** Engraftment of hCD33^+^hCD45^+^ human leukemic cells was assessed after 8-12 weeks (n=4 PDX), and each dot represents the relative engraftment in each mouse. **K-N.** To assess the role of cycling and quiescent cells in AML propagation, PDX cells labeled ex vivo with CFSE were injected into NSG mice, and CFSE retention was measured by flow cytometry after one week. The experimental design is represented schematically (**K**). The CFSE intensity (**L**) and percentage (**M**) were quantified in the quiescent and cycling groups. Post-hoc analysis of CFSE retention was performed in Ki67^-^ and Ki67^+^ subpopulations of leukemic cells from group of mice transplanted with the cycling or quiescent fraction (**N**). The standard deviations are indicated by the vertical bars. The significance levels are denoted by ns (not significant), *p<0.05, **p<0.01, and ***p<0.001.

**Table 1.**
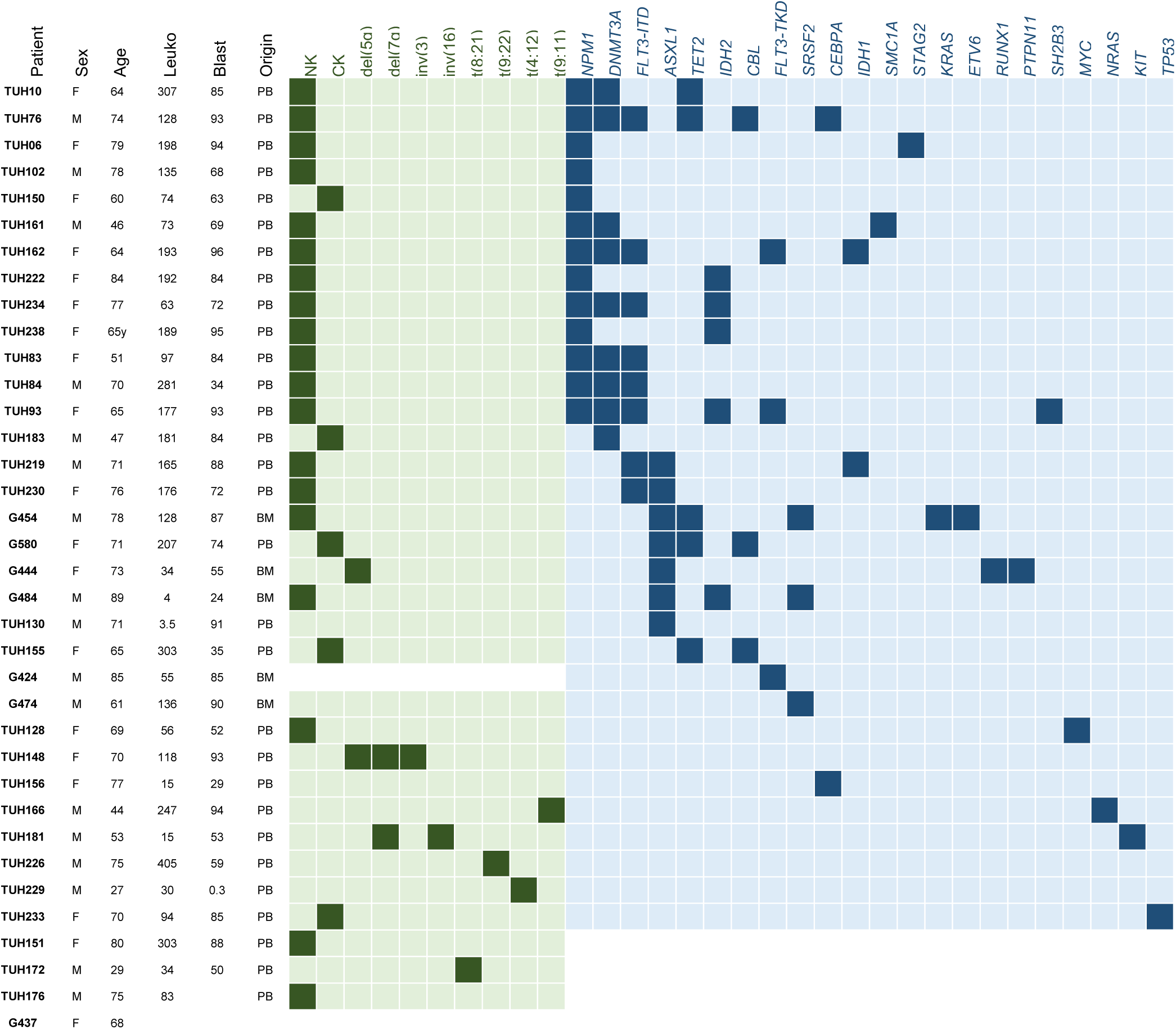
Characteristics of patients with AML. Patients are anonymized and samples are identified using a code in the patient column. Sex: M for male, F for female. Age in years. Leuko: leukocytes in the peripheral blood at the time of sample collection (x10^9^/L). Blast: percentage of bone marrow blast cells. Origin: PB: peripheral blood, BM: bone marrow. Karyotype: NK: normal karyotype, CK: complex karyotype, del: deletion, inv: inversion. Karyotype alteration: green, genetic alteration: blue, no abnormality: grey, analysis not dose: white.

Next, we assessed the in vivo cycling dynamics of human AML cells in PDX models employing labeling assays. In a first set of experiments, we administered Bromodeoxyuridine (BrdU), a synthetic thymidine analogue, to mice for seven consecutive days. We then assessed the level of BrdU incorporation, which serves as a reliable marker of cellular proliferation (*19*), in both quiescent (Ki67^-^) and cycling (Ki67^+^) subsets of AML cells (**Figure 1D-E**). We observed that the proportion of BrdU^+^ cells varied across PDXs, and that BrdU incorporation was higher in Ki67^+^ compared to Ki67^-^ human leukemic cells (**Figure 1F**). However, the tracker labeled both quiescent and cycling subpopulations, indicating that a proportion of quiescent cells had a previous cycling history (**Figure 1F**). To gain a comprehensive understanding of the phenotypic heterogeneity within the quiescent and cycling subsets in vivo, we assessed cell cycle phases and proliferation within the CD34^+^CD38^−^ fraction, which typically represents a small subset that enriched in long-term self-renewing LSCs in the majority of AML cases (*13, 20, 21*) (**Figure S1D**). Our findings revealed that the CD34^+^CD38^-^ cell subpopulation comprised a higher proportion of Ki67^-^ quiescent leukemic cells, in comparison to the CD34^+^CD38^+^ progenitor-like fraction in vivo (**Figure 1G**). Additionally, CD34^+^CD38^-^ LSCs had a lower retention of BrdU labelling compared to CD34^+^CD38^+^ cells in vivo, while the proportion of BrdU^+^ cells was lower in more mature CD34^-^ population of leukemic blasts (**Figure 1H**). The findings identified a critical population of quiescent phenotypically and functionally defined LSCs in vivo.

We further investigated the distinctive characteristics of quiescent and cycling subpopulations through in vivo examination using PDX assays. To achieve this, we initially sorted hCD45^+^hCD33^+^ human AML cells according to their quiescent or cycling phenotype using Pyronin Y (fluorescent RNA-binding dye, does not label G_0_ cells (*22*)), and subsequently xenotransplanted these distinct subpopulations into immunodeficient NSG mice, as reported (*17*) (**Figure 1I**). During the initial stages following transplantation, known as the homing process, we discerned a minimal population of hCD45^+^ cells within the murine bone marrow microenvironment that was quantitatively comparable in mice transplanted with the cycling or the quiescent fraction of leukemic cells (**Figure S1E-F**). In contrast, our results revealed that the quiescent fraction had increased long-term engraftment capacities after 8-12 weeks, thus functionally substantiating an enrichment of leukemia-initiating cells (LICs) in comparison to the cycling subset, as anticipated (*3, 17, 21*) (**Figure 1J**).

In order to monitor cell proliferation during the initial stages of leukemia progression in vivo, we tagged quiescent and cycling fractions ex vivo with carboxyfluorescein succinimidyl ester (CFSE) (*23*) before xenotransplantation (**Figure 1K**). Upon evaluation at the 7-day mark post-transplantation, it was determined that a substantial proportion of human leukemic cells from mice transplanted with the cycling fraction were synchronized and had gone through several cell divisions, as evidenced by their uniform dilution of CFSE (**Figure 1L**). Although cells originating from the quiescent fraction exhibited a higher aggregate level of CFSE retention, their variable CFSE distribution pattern revealed that a noteworthy subset of these cells had undergone active rounds of cell division (**Figure 1L-M**). Using Ki67 staining on day-7 samples, we observed a comparable cell-cycle repartition irrespective of the transplanted fraction (**Figure 1N**). Significantly, a uniform dilution of CFSE was detected within the Ki67^+^ and Ki67^-^ subsets of leukemic cells obtained from mice that underwent transplantation with the cycling fraction (**Figure 1N**). In comparison, CFSE exhibited a similar degree of dilution within the Ki67^+^ leukemic cells obtained from mice that underwent transplantation with the quiescent fraction, while the Ki67^-^ subpopulation from the same group exhibited nearly complete retention of CFSE attesting that these cells had not proliferated in vivo (**Figure 1N**). These results show that quiescent and cycling subsets display distinct proliferation patterns during leukemia propagation in vivo.

Taken together, our findings reveal the presence of a quiescent cell subpopulation characterized by a distinct proliferation pattern and augmented LIC potential when compared to its cycling counterpart.

### Molecular Attributes of Leukemia Stem Cells are Found in the Quiescent Subset of AML Cells

We aimed to identify distinct features of sorted quiescent cells through transcriptomic and single-cell analysis approaches. Through a differential gene expression analysis, we identified 328 and 200 genes that were significantly upregulated in cycling or quiescent subpopulations, respectively (**Figure S2A**). Gene set enrichment analysis on these gene lists uncovered a specific enrichment in LSC-related gene sets among quiescence-related genes (**Figure S2B**). Notably, this included the LSC104 signature derived from an extensive panel of primary AML fractions, which had been functionally evaluated for their in vivo engraftment capabilities in immunodeficient mice (*24*), underscoring the functional relevance of the identified quiescent cellular state (**Figure S2B** and **Table S1**). In addition, the molecular profile of the quiescent subset matched with both quiescent and primed leukemia stem and progenitor cell (LSPC) signatures that were recently uncovered from single-cell profiling of AML samples (*25*) (**Figure S2B** and **Table S1**). These results established a correlation between our molecular profile of quiescence and the functionally defined signatures of AML LSCs. Following an in-depth analysis of the genes upregulated in the quiescent fraction, we curated a concise list of 37 genes that constituted a novel quiescent AML up (Q_AML_UP) signature (**Figure 2A** and **Table S2**). Conversely, we delineated a cycling signature (C_AML_UP) from genes upregulated within the cycling fraction (**Figure 2B** and **Table S2**). Remarkably, we observed that the genes defining the Q_AML_UP signature were absent in both the LSC104 and LSPC quiescent signatures, pointing out the distinctive nature of this novel gene set (**Figure 2B**). These findings show that leukemic cells identified through their quiescence status are reminiscent of LSCs.

**Figure 2.**
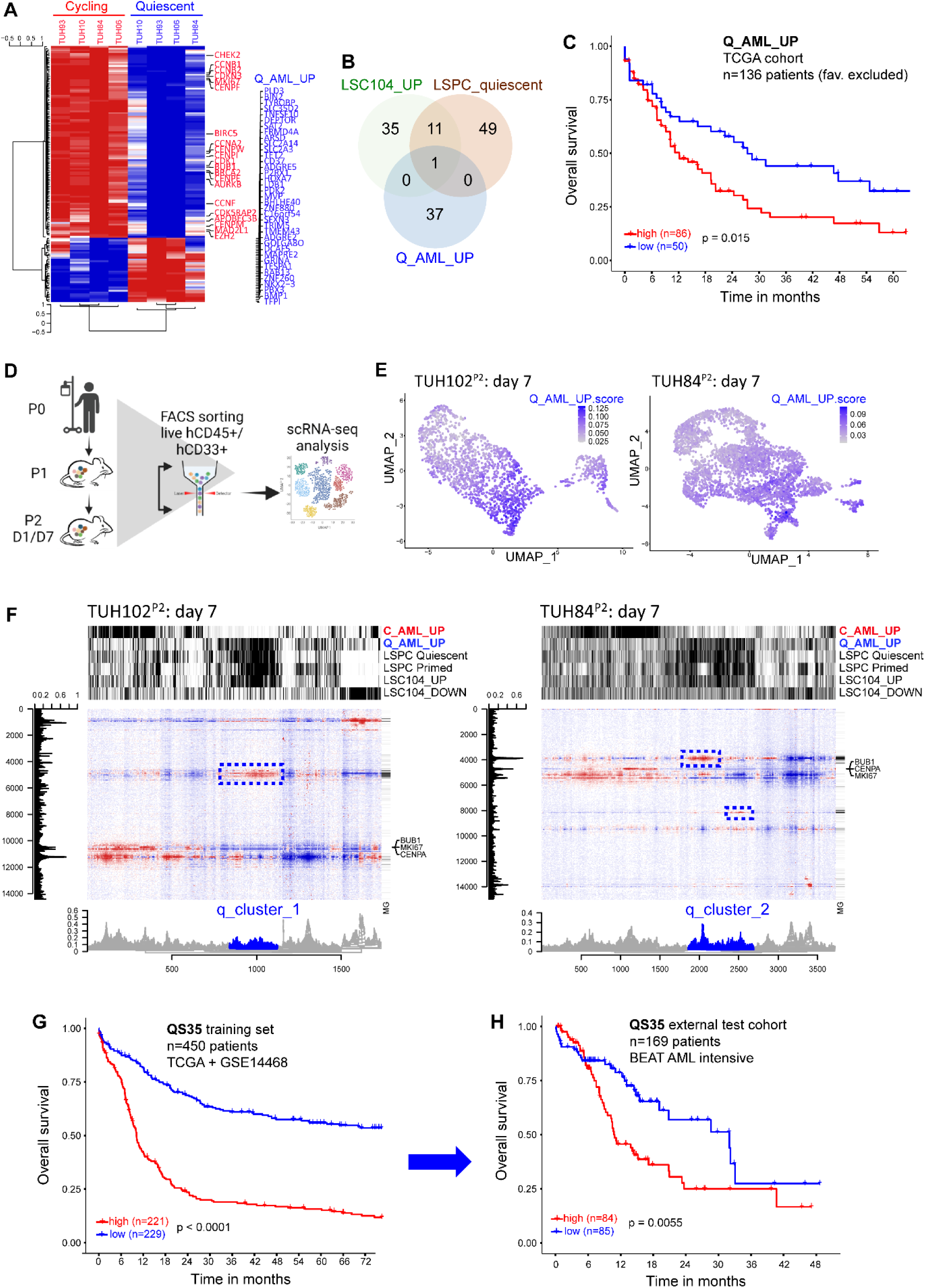
Molecular Attributes of Leukemia Stem Cells are Found in the Quiescent Subset of AML Cells. **A.** Sorting of AML cells into cycling (Pyronin Y^+^) and quiescent (Pyronin Y^-^) populations to perform differential gene expression (DGE) analysis and transcriptomic profiling in four distinct patient-derived xenografts (PDX). The cycling AML up (C_AML_UP) and quiescent AML up (Q_AML_UP) signatures were refined utilizing limma (*69*), and the differential expression of these genes in cycling or quiescent fractions is presented in a heatmap. The names of the 37 genes defining the Q_AML_UP signature is provided, along with several out of the 110 genes defining the C_AML_UP signature. **B.** Venn diagram displaying the overlap between the Q_AML_UP signature and two key molecular signatures associated with leukemic stem cells (LSC104 (*24*) and LSPC quiescent (*25*)). **C.** Evaluation of the impact of the Q_AML_UP signature on overall survival in the TCGA (*26*) AML cohort analyzed following the exclusion of patients classified under the favorable category (n=136 patients). **D.** Primary AML samples (P_0_) were utilized to establish patient-derived xenografts (PDX) denoted as P_1_ samples. Serial transplantation resulted in the generation of P_2_ samples, which were analyzed at one- or seven-days post-transplantation. Human leukemic cells were isolated using CD33/CD45 staining, and their transcriptomes were comprehensively analyzed at the single-cell level utilizing 10X Genomics technology. **E.** Uniform manifold approximation and projection (UMAP) plot for enrichment in the Q_AML_UP signature in the samples TUH102^P2^ and TUH84^P2^ generated using Seurat. **F.** Unsupervised clustering analysis was performed to explore the cellular heterogeneity and gene expression profiles in samples TUH102^P2^ and TUH84^P2^ (day 7). The black and white annotation bars show the value of the Q_AML_UP, C_AML_UP, LSC104, LSPC quiescent, and LSPC primed signatures (White: low, black: high). Cells (Y-axis) are plotted versus genes (X-axis). The gene enriched in the q_cluster_1 and q_cluster_2 from TUH102^P2^ and TUH84^P2^ day 7 samples, respectively that displayed enrichment in both Q_AML_UP and LSC-related signatures are highlighted with blue dash-lined boxes. MG: marked genes. **G-H.** Genes enriched in quiescent LSC cell clusters were used to train a new signature predictive of survival in patients with AML (QS35 signature). **G.** QS35 signature in the training data set (pool of TCGA and GSE14468, n=450 patients). **H.** QS35 signature in the validation cohort constituted of 169 patients from the BEAT AML cohort who were included based on their receipt of an intensive treatment. The p-values determined using the median cut-off are reported.

We evaluated the prognostic implications of the Q_AML_UP signature in AML cohorts by incorporating its gene expression values into patient datasets. In the TCGA dataset (*26*), our investigation revealed that patients exhibiting an enrichment of the Q_AML_UP signature experienced the most unfavorable survival outcome (**Figure 2C**). Interestingly, we found a correlation between gene sets associated with LSCs and the Q_AML_UP signature in the TCGA cohort (**Figure S2C**). However, we did not observe any correlation between these signatures and the LSC17 signature, which was specifically developed as a rapid risk assessment score for newly diagnosed patients with AML (*24*) (**Figure S2C**). As expected, the LSC17 score risk performed well as prognostic marker within the TCGA cohort (**Figure S2D**). Remarkably, the combination of Q_AML_UP and LSC17 signatures enabled the categorization of patients from the TCGA cohort into four groups, with those possessing high values for both gene sets demonstrating the poorest overall survival compared to other groups (**Figure S2E**). These findings highlight the potential of the Q_AML_UP signature as an independent prognostic biomarker in AML.

To characterize quiescent subsets at a single-cell resolution, we employed single-cell RNA sequencing (scRNA-seq). Primary specimens of AML, designated as P_0_ samples, were employed to establish PDXs, denoted as P_1_ samples. The P_1_ samples underwent an iterative transplantation process to generate the P_2_ samples that were then analyzed at seven days post-transplant (**Figure 2D**). Notably, the selection of the 7-day post-transplant timeframe was primarily based on the distinctive cell cycle trajectories revealed by labeling assays for the quiescent and cycling fractions at this specific time point (**Figure 1K-N**). Human leukemic cells were isolated using CD33/CD45 staining, followed by comprehensive analysis of their transcriptomes at the single-cell level (**Figure 2D**). Using the Seurat methodology, we demarcated distinct cell clusters based on shared gene expression patterns (**Figure S2F**). Notably, our analysis identified a specific cluster enriched in Q_AML_UP signature, representing 287 (17%) and 839 (22.5%) cells within the TUH102^P2^ and TUH84^P2^ samples, respectively (**Figure 2E**). Through the application of unsupervised clustering analysis on single-cell transcriptomes, our study unveiled that these unique patterns of cells defined as q_cluster_1 and q_cluster_2 in TUH102^P2^ and TUH84^P2^ samples, respectively harbored a strong and concomitant enrichment in the Q_AML_UP gene expression profile and molecular signatures linked to LSC biology (**Figure 2F**). Notably, we observed that genes upregulated in both q_cluster_1 and q_cluster_2 cell populations (termed q_cluster genes, n=1126) delineated a comparable enrichment pattern than the Q_AML_UP signature (**Figure S2G**). In contrast, we have uncovered a distinctive cellular cluster that exhibits a high expression of the C_AML_UP signature, which highlighted an elevated expression of genes associated with cell proliferation, including *BUB1*, *MKI67* and *CENPA*. Notably, this cluster is devoid of any discernible association with the cell cluster enriched in signatures characteristic of LSCs and/or quiescence (**Figure 2F**). Furthermore, our investigation revealed a share set of 14 genes that were common between the q_cluster genes and the Q_AML_UP signature (**Figure S2H** and **Table S2**). These findings aligned well with our observations in bulk transcriptomes, indicating that both quiescence and LSC-related signatures are comparably rooted in the underlying biology of AML.

We aimed to assess the potential significance of quiescent LSCs characteristics as a prognostic biomarker in AML. We developed a prognostic model for overall survival by analyzing the q_cluster genes in a random selection of 450 cases of AML from the TCGA and GSE14468 cohorts (*26, 27*) (used as the training dataset) with internal cross-validation (**Figure 2G** and **Figure S2H**). The optimal model consisted of a specific combination of 35 genes, collectively known as the quiescence signature 35 (QS35). To assess the validity of the QS35 signature, we employed the remaining 101 patients from the combined TCGA and GSE14468 cohorts, which were not utilized in developing the signature (referred to as the internal dataset), as well as an external dataset comprising 169 patients from the BEAT AML cohort (*28*) (**Figure 2H** and **Figure S2I**). Interestingly, the QS35 signature demonstrated a favorable comparative performance to LSC17 in predicting the risk of mortality within the external validation cohort (**Figure S2J**). Through the joint analysis of bulk and single-cell AML cells transcriptomes, we inferred a pioneering gene signature of prognostic significance in AML patient cohorts.

These findings provide novel insights into the molecular characteristics of quiescent LSCs and their prognostic implications in AML.

### Autophagy-Dependent Iron Pool Sustains the Viability of Quiescent AML Cells

We next aimed to elucidate novel and potentially druggable molecular specificities of quiescent LSCs in AML. To achieve this, we conducted a pathway analysis (*29*) of gene expression patterns in sorted cycling or quiescent fractions. As anticipated, our analysis uncovered an enrichment of cell cycle-related pathways in the cycling fraction (**Figure 3A**). Additionally, the cycling subset demonstrated an enrichment in both translation and mitochondria pathways (**Figure 3A**). To delve deeper into translation regulation, we investigated mammalian target of rapamycin complex 1 (mTORC1) signaling (*30*), and observed higher mTORC1 activity in cycling compared to quiescent leukemic cells (**Figures S3A-D**). Furthermore, cycling cells displayed increased mitochondrial mass and oxidative metabolism, leading to enhanced production of reactive oxygen species (ROS) compared to the quiescent subset (**Figure S3E-G**). On the other side, we identified a distinct set of genes, implicated in lysosome and autophagy pathways, which were enriched within the quiescent fraction of leukemic cells (**Figure 3A**). This finding aligns with the lower activity of mTORC1 observed in quiescent cells, given its physiological role in suppressing autophagy (*31*). To investigate the autophagic flux disparities between quiescent and cycling subtypes of AML cells, we impeded the phagosome-lysosome fusion process by employing Lys05 (*32*), a dimeric analogue of chloroquine, ex vivo in primary AML cells. We observed that Lys05 treatment led to a more pronounced cyto-ID staining, indicating an increased accumulation of phagosomes (*33*) in quiescent AML cells as compared to their cycling counterparts, while lysosomal mass was comparable between these two subpopulations (**Figure 3B** and **Figure S3H**). These results are congruent with transcriptomic analysis and show a marked phagosomal accumulation under lysosome blockage conditions, thereby revealing an intensified autophagic flux in quiescent AML cells compared to their cycling equivalents.

**Figure 3.**
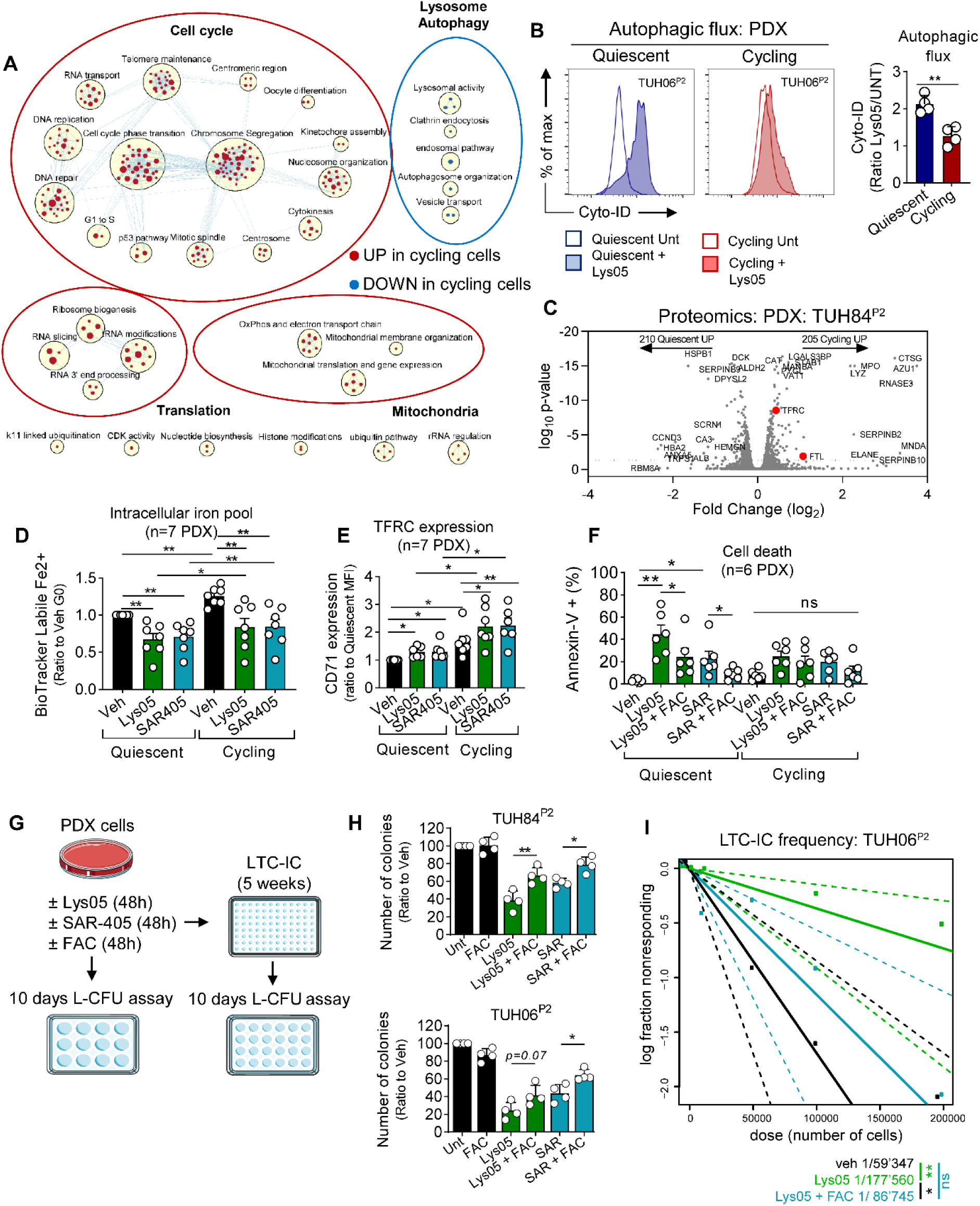
Autophagy-Dependent Iron Pool Sustains the Viability of Quiescent AML Cells. **A.** Transcriptomics analysis was performed on sorted cycling and quiescent leukemic cells using pyronin Y / Hoechst staining. Pathway enrichment analysis through GSEA and visualization using the Cytoscape software’s enrichment map module were conducted. Circles represent pathways, clusters represent biological processes, and lines connect pathways with common genes. Red nodes indicate upregulated pathways, while blue nodes indicate downregulated pathways in cycling compared to quiescent cells (FDR<0.01). **B.** Leukemic cells were incubated with vehicle or 2 μM Lys05 for 12 h, and then stained with the cyto-ID dye. Next, quiescent and cycling subpopulations were distinguished using Ki67 and DAPI. Representative histograms and quantification of the ratio of cyto-ID staining between Lys05 and vehicle conditions are shown (n=4 different PDX). **C.** Quantitative proteomic analysis using LC-ESI-MS/MS was performed on sorted quiescent and cycling fractions separately (n=3 biological replicates from the TUH84^P2^ sample). Results are presented as fold changes versus p-value for the comparison between cycling and quiescent fractions. Proteins involved in iron metabolism with significant differences (FDR ≤ 0.05, log2FC ≥ |0.58|) between the two conditions are highlighted in red. **D-E.** AML cells were incubated with 2 μM Lys05, 5 μM SAR405 or vehicle for 24 hours. Following treatment, cells were subjected to staining using the BioTracker labile Fe2+ dye or anti-CD71 (TFR1) antibody. Quiescent and cycling cell subsets were identified by utilizing Ki67 and DAPI staining. The obtained results are depicted as a ratio relative to the vehicle condition specifically within the quiescent cell subpopulation. Results of intracellular iron pool quantification (n=7) are shown in (**D**), and results of TFR1 membrane expression (n=7) are shown in (**E**). **F-I.** Leukemic cells were incubated ex vivo with 2 μM Lys05, 5 μM SAR405, or vehicle, with or without 10 mg/mL FAC for 48 hours. **F.** Annexin V staining was performed to quantify cell death within quiescent and cycling subpopulations (n=6). **G-I.** The cells were then subjected to L-CFU assays in methylcellulose for 10 days or long-term culture initiating cell assays (LTC-ICs) on MS5 murine stromal cells for 5 weeks followed by secondary seeding on methylcellulose for 10 days. **G.** Schematic representation of the experiments. **H.** Colony formation from two different PDX AML samples was quantified at day 10, and the results are presented as a ratio compared to the vehicle condition. **I.** The frequency of LTC-ICs was quantified using the ELDA software and is indicated for each condition. The standard deviations are indicated by the vertical bars. The significance levels are denoted by ns (not significant), *p<0.05, **p<0.01, and ***p<0.001.

Given the heightened levels of autophagy which bear significant implications for intracellular protein turnover dynamics (*34*), and the lower activity of protein translation components observed in quiescent AML cells, we hypothesized that conducting a comparative analysis of the proteome between quiescent and cycling fractions could potentially reveal biological functions that were not apparent through differential gene expression analysis. We performed a LC/MS-based quantitative protein assessment in both quiescent and cycling subpopulations of AML cells, and observed that out of the 4775 proteins analyzed, 205 and 210 exhibited a statistically significant augmentation in the cycling and quiescent fractions, respectively (**Figure 3C**). Moreover, the biological significance of the respective cell fractions was effectively captured by the differential protein expression, as evidenced by the enrichment of the proteome associated with quiescent cells in molecular signatures characteristic of LSCs (**Figure S3I**). Significantly, our investigation revealed a marked upregulation of pivotal players in iron metabolism, namely transferrin receptor 1 (TFRC) and ferritin light chain (FTL), in actively cycling cells (**Figure 3C**). Therefore, proteomics uncovered changes in iron metabolism pathway between quiescent and cycling leukemic cells that were not captured by transcriptomic analysis.

Considering the function of autophagy in the regulation of iron metabolism and especially of the degradation of iron-containing proteins including TFRC and ferritin (*35*), we investigated the effect of autophagy-dependent iron regulation in both quiescent and cycling subpopulations of AML cells. In tandem with Lys05, we employed the potent vps34 inhibitor SAR405 to block autophagy via the inhibition of nascent autophagosomes assembly (*36*). Notably, these autophagy inhibitors had no impact on the repartition of leukemic cells among G_0_, G_1_ or S-G_2_-M phases of the cell cycle (**Figure S3J**). To explore the dynamics of the intracellular labile iron pool, we employed a dye with selectivity towards Fe^2+^ ions, excluding other metallic species (*12*). Our analysis showed that the cycling AML cells exhibited a markedly expanded intracellular iron pool compared to their quiescent counterparts, and that treatment with autophagy inhibitors effectively depleted the labile iron pool in both cellular subpopulations (**Figure 3D**). In agreement with proteomic analysis, TFRC membrane expression was higher in cycling compared to quiescent AML cells, and treatment with Lys05 or SAR405 caused an increase TFRC expression especially within the cycling subpopulation (**Figure 3E**). Our findings indicate that suppression of autophagic flux decreased iron bioavailability in leukemic cells, along with a compensatory increase in TFRC expression that is less efficient in the quiescent compared to the cycling fractions.

Next, we investigated the impact of autophagy and iron metabolism on the viability of leukemic cells. Short-term assays revealed that Lys05 and SAR405 treatment selectively induced cytotoxicity in quiescent AML cells, with a stronger effect observed for Lys05 (**Figure 3F**). Notably, we found that ferric ammonium citrate (FAC) supplementation could rescue the pro-apoptotic effects of Lys05 and SAR405 in the quiescent cell subpopulation, showing that autophagy-dependent iron availability selectively sustained the viability of quiescent leukemic cells (**Figure 3F**). Leukemia colony-forming units (L-CFU) and leukemic long-term culture-initiating cells (LTC-IC) assays serve as valuable ex vivo surrogates for assessing the functionality of progenitor leukemic cells and LICs (*37–39*). We evaluated the efficacy of Lys05, SAR405, or their combination with FAC on primary AML cells using short-term L-CFU assays on methylcellulose and longer-term LTC-IC assays on murine stromal cells coupled to secondary plating in methylcellulose (*40*) (**Figure 3G**). We observed that treatment with autophagy inhibitors significantly decreased colony formation in L-CFU assays after 10 days compared to the control condition, and this effect was partially rescued by co-incubation with FAC (**Figure 3H**). Moreover, Lys05 treatment reduced the frequency of LTC-ICs compared to the control, and this effect was attenuated by incubation with FAC (**Figure 3I**). These results show that autophagy inhibition depleted progenitors and LSCs ex vivo dependent on iron metabolism.

Taken together, the present findings show that autophagy-dependent iron bioavailability is a critical component of the subpopulation of quiescent leukemic cells, and is required for the proper function of progenitor and LSCs in AML.

### Low Iron Metabolism is Linked to Quiescent Leukemic Stem Cells in Primary AML Cells

Primary specimens from patients with AML represent the original source of disease and therefore provide a more clinically relevant context compared to leukemia cells originating from PDXs (*6, 41*). Consequently, we conducted an in-depth analysis of a cohort comprising 36 primary AML samples, which were utilized for a range of ex vivo assays (**Figure 4A** and **Table 1**). Notably, the cytogenetic and molecular profiles observed in these patients displayed the characteristic hallmark heterogeneity associated with AML (*26, 42*) (**Figure 4B** and **Table 1**). We performed a comprehensive cell cycle analysis of primary AML samples and defined four distinct subpopulations of leukemic cells based on CD34 and CD38 staining (**Figure 4C**). While a majority of primary leukemic cells from patients with AML were quiescent, as measured by negative staining for both Ki67 and propidium iodine (PI), our results revealed that the CD34^+^CD38^-^ fraction displayed a significantly higher proportion of Ki67^-^PI^-^ quiescent cells compared to the progenitor-like CD34^+^CD38^+^ immature leukemic cells (**Figure 4C-D**). Here, we therefore established a correlation between LSC phenotype and a state of quiescence in primary AML cells, consistent with findings from PDX models. Furthermore, we observed a distinct iron metabolism profile in CD34^+^CD38^-^ LSCs characterized by lower labile iron pool, reduced membrane expression of TFR1, and reduced capacities for transferrin uptake relative to CD34^+^CD38^+^ progenitor-like cells, and also to more mature CD34^-^ subpopulations (**Figure 4E-G**, **Figure S4A-C** and **Table S3**). These findings indicate that the reduced iron uptake observed in quiescent AML cells from PDX assays is recapitulated in phenotypically defined CD34^+^CD38^-^ LSCs.

**Figure 4.**
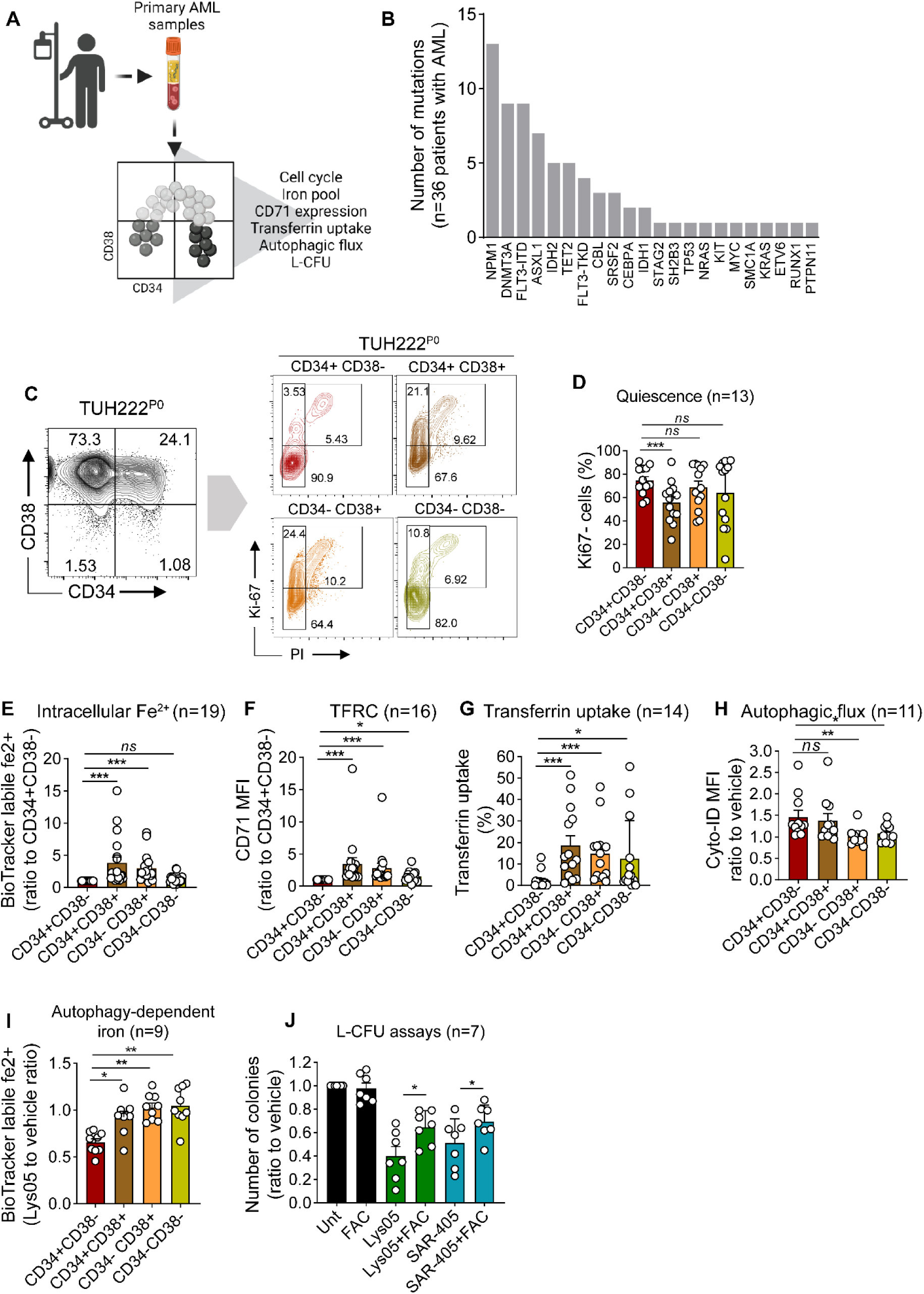
Low Iron Metabolism is Linked to Quiescent Leukemic Stem Cells in Primary AML Cells. **A.** Primary leukemic cells were isolated and characterized from peripheral blood or bone marrow samples obtained from patients diagnosed with AML. Mononuclear cells were enriched via gradient centrifugation and cryopreserved. Upon thawing, leukemic cells were identified using hCD45 versus SSC-A and subsequently stained with anti-CD34 and CD38 antibodies for classification into four distinct groups based on these markers. **B.** Genomic alterations in driver genes implicated in AML were investigated within a cohort consisting of 36 AML patients. **C.** Representative contour plots illustrate CD34, CD38, Ki67, and propidium iodine (PI) staining in a primary AML sample. The left panel demonstrates CD34/CD38 gating, while the right panel presents Ki67/PI gating within each of the four subpopulations. **D-G.** Quantitative analysis of various parameters within each of the four leukemic cell subpopulations defined by CD34/CD38 markers. **D.** Proportion of quiescent Ki67^-^ cells (n=13). **E.** Intracellular Fe^2+^ quantification using the BioTracker Labile Fe^2+^ dye (n=19). **F.** TFR1 expression quantification at the cell surface (n=16). **G.** Measurement of transferrin uptake using AF488-labelled transferrin (n=14). **H-I.** Leukemic cells were incubated with vehicle or 2μM Lys05 for 12 h, and the four leukemic cell subpopulations defined by CD34/CD38 markers were analyzed. **H.** Autophagic flux quantification using cyto-ID staining (n=11). **I.** Intracellular Fe^2+^ quantification using the BioTracker Labile Fe^2+^ dye (n=9). **J.** Leukemic cells were incubated ex vivo with 2 μM Lys05 or 5 μM SAR405 or vehicle, and/or 10 mg/mL FAC for 48 hours. Subsequently, cells were seeded in methylcellulose for 10 days in L-CFU assays. Colony formation was quantified at day 10, and the results are presented as a ratio to the vehicle condition (n=7). Standard deviations are indicated by vertical bars. Significance levels are denoted as ns (not significant), *p<0.05, **p<0.01, and ***p<0.001.

Next, we aimed to explore the role of autophagy-dependent iron metabolism in primary AML specimens. Firstly, we examined the autophagic flux induced by Lys05 in different subpopulations of CD34^+^ and CD34^-^ cells. Our data revealed a significant accumulation of phagosomes in CD34^+^ compared to CD34^-^ cells upon Lys05 treatment, albeit results were comparable between CD34^+^CD38^-^ and CD34^+^CD38^+^ fractions (**Figure 4H** and **Figure 4SD**). However, treatment with Lys05 resulted in a substantial decline in the labile iron pool in CD34^+^CD38^-^ LSCs compared to other sub-fractions and especially CD34^+^CD38^+^ cells (**Figure 4I**). These results showed that maintenance of the pool of intracellular iron was more reliant on autophagy in CD34^+^CD38^-^ LSCs compared to other AML subpopulations. Additionally, we performed L-CFU assays to evaluate the effect of autophagy inhibition on colony formation in primary AML cells. Our results showed that treatment with either Lys05 or SAR405 significantly reduced colony formation, which was blunted by exogenous iron supply (**Figure 4J**). These results confirmed that autophagy-dependent iron pool was a critical dependency of LSCs in primary AML specimens.

Our findings offer compelling evidence that in the context of AML patients, iron-dependent autophagy represents a specific vulnerability of LSCs.

### Ferritinophagy Inhibition via NCOA4 Impairs Leukemia Stem Cells Function in AML

Ferritinophagy is a specialized form of autophagy that requires the cargo receptor protein NCOA4 to selectively target and deliver ferritin to lysosomes for degradation (*10*). We investigated the role of NCOA4, and observed that NCOA4 expression was significantly higher in quiescent compared to cycling AML cells (**Figure S5A**). Moreover, NCOA4 top co-dependencies involved iron metabolism and autophagy effectors in multiple cancer cell lines (**Figure S5B**). To explore the functional significance of this observation, we employed two different mCherry-tagged shRNAs to specifically target NCOA4 expression in AML cells sourced from PDXs (**Figure 5A** and **Figure S5C**). Our findings indicate that NCOA4 depletion resulted in a significant reduction of the labile iron pool in both quiescent and cycling AML cells subpopulations (**Figure S5D-E**). Next, we conducted NCOA4 depletion experiments in two human AML cell lines, namely MOLM-14 and OCI-AML2 (**Figure S5F**). Remarkably, we observed that NCOA4 knockdown led to a decrease in the in vitro growth of both cell lines, which could be reversed by exogenous iron supplementation (**Figure S5G**). These findings underscore the critical role of NCOA4 in regulating iron metabolism and the viability of AML cells.

**Figure 5.**
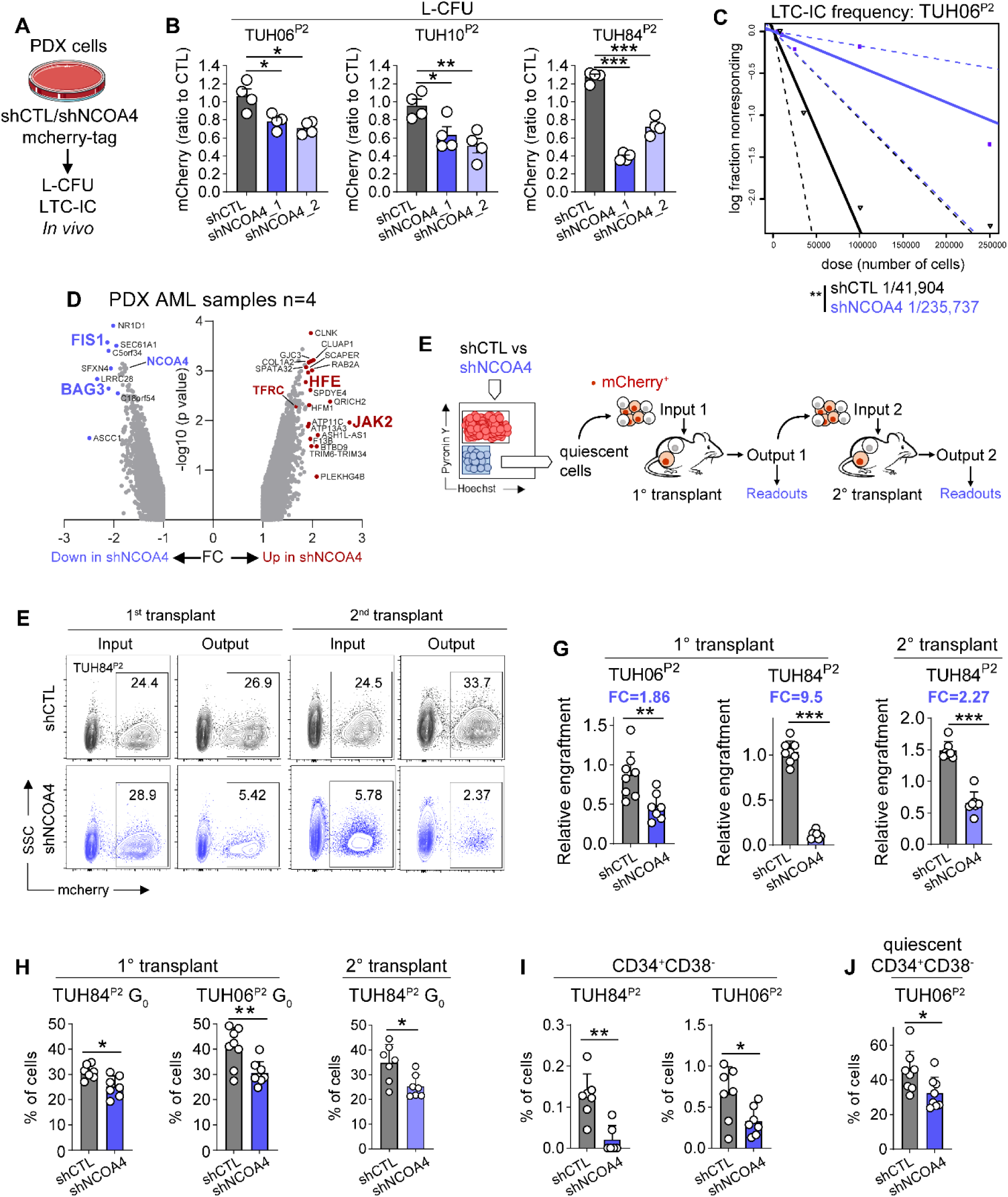
Ferritinophagy Inhibition via NCOA4 Impairs Leukemia Stem Cells Function in AML. **A-J.** AML cells were transduced ex vivo with lentiviral vectors expressing mCherry-tagged NCOA4-targeting or control shRNAs, followed by various in vitro and in vivo experiments. **A.** Schematic representation of the experimental design. **B.** L-CFU assays were conducted in leukemic cells transduced with control or two different anti-NCOA4 shRNAs (shNCOA4_1 and shNCOA4_2). Results from biological replicates (n=4) are presented as a ratio to the control condition (n=3 different PDXs). **C.** LTC-IC assays were performed in leukemic samples transduced with control or shNCOA4_1 shRNAs. The frequency of LTC-ICs was quantified using the ELDA software. The calculated frequencies of LTC-IC are provided for each experimental group. **D.** Differential gene expression analysis in NCOA4-depleted leukemic cells relative to the control condition (n=4). Genes of interest among the top-modulated differentially expressed genes are highlighted. **E-J.** AML cells transduced with control or shNCOA4_1 shRNAs were sorted based on Pyronin-Y/Hoechst staining, and the quiescent fraction was transplanted into immunodeficient NSG mice (n=2 different PDX assays, n=7-8 mice per experimental group). Following leukemia propagation in primary recipient mice, subsequent xenotransplantation was conducted to investigate the self-renewal functions of LSCs. **E.** The proportion of efficiently transduced mCherry^+^ cells before transplant represented the input. After 10-12 weeks in vivo, the engraftment of mCherry^+^ cells defined the output. **F.** Representative flow cytometry histograms. **G.** Relative engraftment defined as the ratio of output to input. Fold-changes (FC) of relative engraftment are provided. **H.** Percentage of human AML cells in the G_0_ phase of the cell cycle, as determined by Ki67/DAPI staining. **I.** Proportion of CD34^+^CD38^-^ cells. **J.** Proportion of quiescent (G_0_) cells among the CD34^+^CD38^-^ subpopulation. The standard deviations are indicated by the vertical bars. The significance levels are denoted by *p<0.05, **p<0.01, and ***p<0.001.

Furthermore, NCOA4 depletion significantly decreased colony formation in L-CFU assays and reduced the frequency of LTC-ICs compared to the control condition (**Figure 5B-C**). Next, we investigated the differential gene expression between control and NCOA4-depleted leukemic cells. Notably, the depletion of *NCOA4* resulted in an increased expression of genes associated with iron uptake, including *TFRC* and its iron sensing partner, *HFE* (*9*) (**Figure 5D** and **Figure S5H**). Moreover, NCOA4-depleted cells displayed an upregulation of *JAK2*, which encodes a tyrosine kinase known to be involved in the production of hepcidin (*9*) (**Figure 5D**). We further observed that the genes *FIS1* and *BAG3* displayed significant downregulation in the NCOA4-depleted leukemic cells (**Figure 5D**). These genes have been known to facilitate mitophagy, and their targeted invalidation has been associated with compromised functions in LSCs and HSCs in the case of *FIS1* or *BAG3*, respectively (*43–45*). Consistent with this finding, a depletion of the Q_AML_UP signature was observed in shNCOA4 leukemic cells compared to their control counterparts (**Figure S5I**). The results of these ex vivo assays suggest that targeting NCOA4, a key effector of ferritinophagy, could serve as a promising therapeutic strategy to selectively eliminate LSCs in AML.

To further examine the role of NCOA4 in the function of LSCs, we performed in vivo experiments in PDX AML models. First, we transduced AML cells ex vivo with mCherry-tagged anti-NCOA4 or control shRNAs using lentivirus. Then, we sorted quiescent leukemic cells based on their low Pyronin-Y staining to perform a first transplantation of this fraction into NSG mice. Following the propagation of leukemia in primary recipient mice, a subsequent xenotransplantation was performed to evaluate the self-renewal functions of LSCs (*18, 46*). The proportion of mCherry^+^ human AML cells was assessed both before (input) and after each round of transplantation (output), enabling the determination of relative engraftment as the ratio of output to input, as reported (*46, 47*) (**Figure 5E**). Remarkably, we uncovered that the depletion of NCOA4 had a notable impact on reducing leukemia propagation from the sorted quiescent leukemic cells within primary recipient mice but also in bulk leukemia samples transplanted into secondary recipient mice (**Figure 5F-G**). Additionally, leukemia propagated from NCOA4-depleted AML cells showed a significant reduction in the quiescent G_0_-phase leukemic cells both after primary and secondary transplantation, with heterogeneous impact on the repartition of G_1_ and S-G_2_-M cells (**Figure 5H** and **Figure S5J-K**). However, NCOA4 depletion did not result in modifications of the differentiation marks CD11b, CD14 or CD15 compared to the control condition (**Figure S5L**). Importantly, we observed a more pronounced decrease in the abundance of leukemic cells exhibiting CD34^+^CD38^-^ LSCs phenotype upon NCOA4 depletion, and also a reduction in CD34^+^CD38^+^ progenitor-like cells, while concurrently an increase in mature CD34^-^ cells were observed (**Figure 5I** and **Figure S5M**). Moreover, depletion of NCOA4 led to a reduction in the percentage of quiescent CD34^+^CD38^-^ LSCs relative to the control group (**Figure 5J**). These findings provide compelling evidence that the key ferritinophagy effector NCOA4 is an essential dependency of LCSs in AML.

Collectively, our observations show that targeting ferritinophagy via NCOA4 depletion impede LSCs functions in AML.

### Ferritinophagy Disruption as a Novel Pharmacological Approach against Leukemic Stem Cells

In order to facilitate the translation of our preclinical findings into clinical applications for AML, we leveraged a recently synthesized small compound inhibitor targeting NCOA4. This inhibitor, named compound **9a**, exerts a direct binding effect on NCOA4, consequently inhibiting its interaction with ferritin heavy chain (FTH1) and leading to the suppression of ferritinophagy (*48*).

First, we performed L-CFU assays from primary AML specimens, and our results showed that compound **9a** decreased the formation of colony after 10 days, specifically at the concentration of 1 μM or 5 μM (**Figure 6A**). Remarkably, when the same concentrations were applied, compound **9a** demonstrated negligible effects on the generation of colony forming unit-granulocyte-macrophage (CFU-GM) and burst forming unit-erythroid (BFU-E) from normal CD34^+^ hematopoietic progenitor cells cultured in methylcellulose during 14 days (**Figure 6B**). Additionally, compound **9a** moderately attenuated the growth rate of normal hematopoietic cells ex vivo at a concentration of 5 μM, while exhibiting no such impact at 1 μM (**Figure S6A**). These observations suggest that the compound displayed efficacy against leukemic cells while exerting minimal influence on normal human hematopoietic progenitor cells at the concentration of 1 µM, which subsequently guided its application in further ex vivo assays.

**Figure 6.**
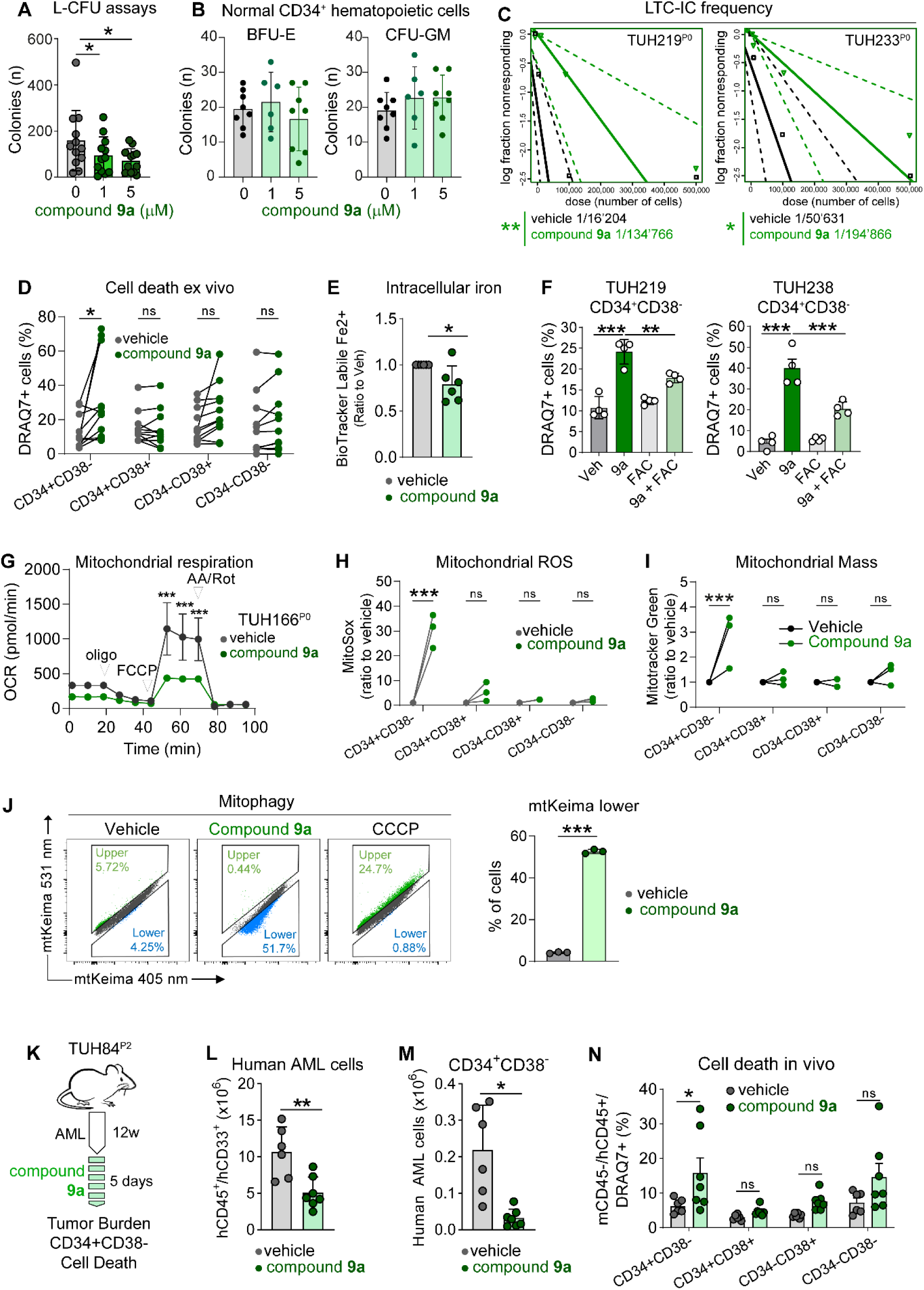
Ferritinophagy Disruption as a Novel Pharmacological Approach against Leukemic Stem Cells. **A.** Primary leukemic cells were incubated with vehicle or 1 or 5 μM compound **9a** and seeded in methylcellulose culture during 10 days (n=12). **B.** Normal CD34^+^ hematopoietic progenitor cells were incubated with vehicle or 1 or 5 µM compound **9a** in methylcellulose cultures. BFU-E and CFU-GM were counted under an inverted microscope at day 14**. C.** LTC-IC assays were performed on leukemic samples cultured in liquid conditions and subjected to either vehicle or 1 µM compound **9a** treatment for 72 hours. Next, the samples were cultured on MS-5 stromal cells for a period of 5 weeks, and subsequently seeded in methylcellulose for 10 days. The frequency of LTC-ICs was quantified using the ELDA software. **D.** Primary AML cells were incubated with vehicle or 1 μM compound **9a** in liquid culture for 24 h (n=11). The proportion of DRAQ7^+^ dead cells was determined within each of the four subpopulations defined by CD34/CD38 staining. **E.** Primary AML cells were incubated with vehicle or 1 μM compound **9a** for 6 h, and intracellular Fe^2+^ was quantified using the BioTracker Labile Fe^2+^ dye (n=6). **F.** Primary AML cells were incubated with vehicle or 1 μM compound **9a** for 48 h, without or with 0.1 mg/mL FAC. The proportion of DRAQ7^+^ dead cells was determined within the CD34^+^CD38^-^ subset (n=2 different primary AML samples, n=4 biological replicates). **G-I.** Primary AML cells were cultured with vehicle or 1 µM compound **9a** for 48 h. **G.** Bioenergetic assays were conducted to measure the oxygen consumption rate (OCR) in mpH/min. Oligomycin (Oligo, mitochondrial ATP synthase inhibitor), Carbonyl cyanide-4 (trifluoromethoxy) phenylhydrazone (FCCP, mitochondrial uncoupling agent), and rotenone and antimycin (AA/Rot, complex I and complex III inhibitors, respectively) were used as specific inhibitors of mitochondrial respiratory chain during the assays. **H-I.** Mitochondrial ROS production and mitochondrial mass were measured within the four subpopulations defined by the CD34 and CD38 markers using MitoSOX and Mitotracker green staining, respectively (n=3). **J.** MOLM-14 cells were transduced with a lentiviral vector allowing the constitutive expression of the mt-keima probe. Then, cells were incubated with vehicle, 1 μM compound **9a** or 1 µM of carbonyl cyanide m-chlorophenylhydrazone (CCCP, a mitochondrial oxidative phosphorylation uncoupler, a positive control for mitophagy induction) for 6 h. Then, fluorescence at 405 nm and 561 nm was measured, and the proportion of cells having a low 561 nm signal is quantified (right panel). **K-N.** Leukemic cells were propagated in NSG mice for 12 weeks. Subsequently, mice received either placebo or 10 mg/kg compound **9a** via daily intraperitoneal injection for five consecutive days. The schematic representation of the experimental setup is shown (**K**) **L.** Burden of human CD45^+^CD33^+^ leukemic cells. **M.** Proportion of CD34^+^CD38^-^ leukemic cells. **N.** Measurement of cell death using DRAQ7^+^ staining within each of the four populations defined by CD34/CD38 markers. The standard deviations are indicated by the vertical bars. The significance levels are denoted by *p<0.05, **p<0.01, and ***p<0.001.

In ex vivo assays, the administration of compound **9a** exhibited a remarkable decrease in the frequency of LTC-ICs in two distinct primary AML samples when compared to the control placebo (**Figure 6C**). This noteworthy impact on the most primitive leukemic cell populations in ex vivo experiments was consistent with our finding that compound **9a** specifically induced cell death within the phenotypically defined CD34^+^CD38^-^ LSC subpopulation (**Figure 6D**). Focusing on iron metabolism, we found that compound **9a** decreased the intracellular iron pool in primary AML cells compared to the vehicle, as anticipated from other cellular models (*48*) (**Figure 6E**). Accordingly, differential gene expression (DGE) analysis revealed that compound **9a** induced changes in iron metabolism-related genes including *TFR1*, *SLC40A1*, *STEAP2* and *ACO1* (**Figure S6B**). In addition, we observed that treatment with compound **9a** led to a depletion in the quiescence molecular signature (Q_AML_UP) compared to the vehicle condition (**Figure S6C**). Remarkably, the elective impairment in cell viability induced by compound **9a** within the CD34^+^CD38^-^ subset was significantly rescued by exogenous iron (**Figure 6F**). These results show that compound **9a** exhibits significant activity against immature leukemic cells ex vivo dependent on the depletion of intracellular iron pool.

From a mechanistic standpoint, we established a correlation between the cytotoxic impact of compound **9a** and the perturbation of mitochondrial oxidative metabolism. This was attested by a reduction in mitochondrial respiration and by a burst in the generation of mitochondrial ROS that were specifically within the CD34^+^CD38^-^ subset (**Figure 6G-H** and **Figure S6D**). Furthermore, treatment with compound **9a** resulted in an increased mitochondrial content within the LSC compartment, suggesting that inhibition of NCOA4 resulted in an accumulation of defective mitochondria (**Figure 6I**). As we showed that NCOA4 controlled *FIS1* and *BAG3* expression in AML cells, we hypothesized that the mitochondrial phenotype induced by compound **9a** could interfere with the mitophagy process. To investigate this possibility, we employed a pH-sensitive fluorescence assay featuring the coral-derived protein Keima, localized within the mitochondrial matrix of the MOLM-14 cell line using an mt-keima probe (ref). Briefly, Keima demonstrates a dual excitation spectrum, manifesting as green fluorescence (peak at 440 nm) under physiological pH conditions (pH 8, typical of mitochondria) and transitioning to a red emission (peak at 586 nm) under acidic conditions. During mitophagy, mitochondrial delivery to lysosomes (pH 4.5) is detected by a characteristic shift in the excitation spectrum of mt-keima (*49*). Notably, treatment of mt-keima-expressing MOLM-14 cells with compound **9a** resulted in a reduced proportion of cells displaying red fluorescence indicative of a low pH environment, as compared to the vehicle-treated condition (**Figure 6J**). These results show that the inhibition of NCOA4 by compound **9a** resulted in an inhibition of mitophagy in MOLM-14 AML cells.

Next, we performed PDX assays where mice were subjected to either placebo or compound **9a** treatment for a continuous duration of five days (**Figure 6K**). The administration of compound **9a** during a limited timeframe decreased the intracellular iron pool and demonstrated a remarkable activity against leukemia in vivo compared to the placebo, leading to a substantial 50% reduction in the overall burden of human AML cells, especially within the CD34^+^CD38^-^ subset (**Figure 6L-M** and **Figure S6E**). As observed after NCOA4 depletion, treatment with compound **9a** resulted in a decreased number of CD34^+^CD38^+^ progenitor-like leukemic cells compared to the placebo. However, the effect on the more mature CD34^-^ subpopulation was more limited and non-significant (**Figure S6F-G**). Additionally, compound **9a** demonstrated the ability to selectively induce cell death within the CD34^+^CD38^-^ population in vivo (**Figure 6N** and **Figure S6H**). Therefore, the anti-leukemic effects induced by compound **9a** in vivo were reminiscent of those resulting from genetic NCOA4 depletion in PDX AML models and particularly targeted the LSC subpopulation. Remarkably, no signs of compound **9a**-induced toxicity were observed throughout the course of the study, as indicated by the stable body weight of the mice, and the comparable amounts and viability of murine CD45^+^ hematopoietic cells found in both experimental groups (**Figure S6I-K**). These results provide evidence of the therapeutic potential of pharmacological NCOA4 blocking in preclinical models of AML.

Within the scope of this study, we present a novel concept that highlights the dependence of quiescent LSCs on ferritinophagy to maintain their intracellular iron pool, as opposed to their more differentiated leukemic counterparts. Therefore, inhibition of ferritinophagy through genetic or pharmacological interference with NCOA4 impairs LSC functions, culminating in the induction of anti-leukemic effects within preclinical models of AML. Thus, ferritinophagy inhibition represents an innovative therapeutic approach for AML patients.

## DISCUSSION

Leukemia stem cells (LSCs) are a rare subset of the leukemic blasts that can self-renew and differentiate into more mature leukemic cells. They are defined based on their ability to initiate leukemia development and maintain the disease in vivo, as well as to recapitulate the phenotype of the original patient’s leukemia when transplanted into immunocompromised mice (*3*). The pool of AML LSCs is predominantly found, albeit not entirely, within the CD34^+^CD38^-^ phenotype that is also linked to normal HSCs (*13, 20, 50*). Several studies have shown that LSCs are present in AML patients before and after chemotherapy, and that the persistence of LSCs after treatment is associated with a higher risk of relapse (*6, 41, 51*). LSCs share many fundamental characteristics with their normal HSCs counterparts, including self-renewal capacity, multipotency, and reliance on similar molecular signaling pathways for survival and differentiation (*52*). Notably, the quiescence of LSCs underlie resistance to cell cycle-dependent chemotherapies in AML (*5, 17, 53–55*). Considerable efforts have been made to explore the role of the bone marrow microenvironment in inducing quiescence in LSCs (*52*). However, the cell-intrinsic molecular mechanisms underlying the quiescent state of LSCs and their role in AML pathogenesis are not fully understood.

In this study, we aimed to capture novel features of quiescent cells in PDX models and primary leukemic cells from patients with AML to identify potential vulnerabilities that could be exploited for therapeutic benefit. We observed that the cell-cycle distribution of leukemic cells in PDX models was similar to that in patients with AML, with only a minority of cells undergoing divisions (*15*). Further experiments revealed that the quiescent fraction displayed a more heterogeneous proliferative trajectory compared to its cycling homolog, with a subset of quiescent cells that did not proliferate during in vivo propagation of leukemia. However, the quiescent fraction showed increased LIC function in vivo, as anticipated (*17*). This convergence of quiescent and LSC profiles was further substantiated through single-cell analysis conducted in vivo, showing that LSCs and quiescence molecular signatures converged within the same cluster of cells. Notably, our findings corroborate earlier investigations that have reported the existence of a quiescent leukemic cell subset demonstrating extensive self-renewal capacities (*55*). Intriguingly, our study also uncovered a novel set of genes – named the QS35 risk score – associated with quiescence that proved to be a robust predictor of poor outcomes in patients with AML, reminiscent of previous studies regarding the prognostic role of LSC abundance and activity (*6, 24, 41, 56–58*).

The similarities between quiescent cells and LSCs observed in our models prompted us to adopt the quiescence perspective for uncovering novel vulnerabilities inherent to LSCs. Our investigation revealed that quiescent cells exhibited reduced mitochondrial and metabolic activity, including decreased production of reactive oxygen species (ROS), compared to their cycling counterparts. Notably, both normal HSCs and LSCs are found enriched in a cell subset defined by their low oxidative metabolic state, known as ROS-low (*59, 60*), suggesting that the functions of quiescent and ROS-low LSCs may largely overlap. In addition, the quiescent subpopulation displayed increased lysosomal and autophagic activity relative to their cycling counterparts. This finding was substantiated by the low activity of the mTORC1 signaling pathway, known to impede autophagy (*31*), found in quiescent leukemic cells. As a possible mechanistic explanation of this result, we showed a substantial upregulation of DEPTOR, a negative regulator of mTORC1 (*30*), within the quiescent subpopulation. These results were reminiscent of a recent study in HSCs showing that transcription factor EB (TFEB) promotes quiescence and self-renewal through induction of the endolysosomal pathway (*61*). Interestingly, our study uncovered that inhibition of autophagy selectively induced cytotoxic effects on the quiescent subpopulation of AML cells. This observation is particularly significant considering the close proximity between the quiescent and LSC subpopulations observed in our study. These findings align with compelling evidence highlighting the role of autophagy in the maintenance of both HSC and LSC populations (*43, 62*).

Through an extensive comparative analysis using proteomics, we have identified iron metabolism as a distinctive hallmark that set apart quiescent and cycling fractions. Specifically, we observed the lowest intracellular iron levels in the quiescent and phenotypically defined CD34^+^CD38^-^ LSC populations, which correlated with reduced membrane expression of TFRC and impaired transferrin uptake in these cells. Furthermore, we discovered that inhibiting autophagy led to a depletion of intracellular iron, while conversely, supplementation of exogenous iron rescued the lethal phenotype observed in quiescent leukemic cells upon autophagy inhibition. Recent investigations have revealed a compelling association between the lysosomal compartment and iron metabolism. Notably, lysosomes have emerged as crucial regulators of intracellular iron distribution. Inhibition of lysosomal acidification has been found to diminish mitochondrial iron availability, thereby impairing mitochondrial functions and triggering cell death (*63*). Furthermore, the retention of iron within lysosomes induced by a small compound irinomycin disrupted iron-dependent mitochondrial metabolism and led to non-apoptotic cell death in AML (*64*). Therefore, we could hypothesize that iron depletion plays a substantial role in the cell death observed upon autophagy inhibition specifically within the quiescent LSC compartment of AML cells.

The preferential susceptibility of the quiescent subset to autophagy-regulated iron pool indicated their reliance on ferritinophagy to fulfill their intracellular iron demand. Therefore, we focused on NCOA4, the main cargo receptor mediating ferritinophagy as potential vulnerability of quiescent LSCs. Our study yielded compelling insights, demonstrating that the inhibition of NCOA4 exhibited significant anti-leukemic effects ex vivo and in vivo, and selectively targeted quiescent and self-renewing LSCs. Notably, we observed an elevated expression of NCOA4 in quiescent leukemic cells when compared to their cycling counterparts, potentially elucidating the selective impact of NCOA4 depletion within the LSC compartment. Moreover, we observed that the mitochondrial fission factor FIS1 emerged as one of the most significantly downregulated genes upon NCOA4 depletion. Notably, FIS1-dependent mitophagy, by ensuring the elimination of damaged mitochondria in AML, serves to maintain the LSC population (*43*). Remarkably, NCOA4 inhibition mediated by compound **9a** resulted in a marked reduction of mitophagy in leukemic cells. This novel function of NCOA4 may involve FIS1 and contribute significantly to the maintenance of the LSC pool.

From a translational standpoint, we showed for the first time the efficacy of the small molecular inhibitor compound **9a** that disrupts the interaction between NCOA4 and ferritin (*48*) in selectively targeting LSCs. This compound exhibited notable anti-leukemic effects both ex vivo and in vivo, as demonstrated in PDX models of AML. Similar to the effects observed upon NCOA4 depletion, the anti-leukemic activity triggered by compound **9a** was rescued with the addition of exogenous iron, substantiating the molecular specificity of this small molecule as a ferritinophagy inhibitor. Specifically, compound **9a** exhibited selective toxicity towards the phenotypically defined CD34^+^CD38^-^ LSC compartment, which was associated with an important production of mitochondrial ROS within the same cellular subset. These findings align with the concept that the preferential utilization of iron, sourced from NCOA4-mediated ferritin degradation, plays a pivotal role in maintaining mitochondrial functionality (*65*). Similarly, a recent investigation revealed the dependence of pancreatic cancer cells on NCOA4-mediated ferritinophagy. In this specific context, ferritinophagy serves as a vital source of endogenous iron, which is utilized for the synthesis of iron-sulfur cluster proteins that play a crucial role in governing mitochondrial oxidative metabolism. Consequently, targeting NCOA4 could hold promise as a potential therapeutic approach for this type of cancer (*12*). These findings highlight the promise of ferritinophagy inhibition as a tailored treatment approach for cancer and AML patients.

In conclusion, our study unveils novel molecular insights into the cell-intrinsic properties of quiescent LSCs, shedding light on the promising therapeutic potential of NCOA4 inhibitors as a groundbreaking approach in the management of AML.

## MATERIAL AND METHODS

### Patients

Primary cells derived from de-identified patients diagnosed with acute myeloid leukemia (AML) were obtained from the HIMIP collection (BB-0033-00060). The HIMIP collection has been duly declared to the Ministry of Higher Education and Research (DC 2008-307, collection 1). Prior to sample acquisition, a transfer agreement (AC 2008-129) was obtained following approval by an ethical committee. In compliance with ethical guidelines and the principles outlined in the Declaration of Helsinki, written informed consent was obtained from all participating patients. The consent granted permission for the utilization of their samples for research purposes. **Table 1** provides a comprehensive summary of the characteristics of the patients included in this study, while ensuring the anonymity of their identities.

### Animals

All animal experiments described in this study adhered to the guidelines set forth by the Association for Assessment and Accreditation of Laboratory Animal Care International. The protocols and procedures were conducted with the approval of the local ethics committee, specifically the Geneva health department, under the authorization GE/123/19 and GE/166. For the experiments involving animal models, 6-8 weeks old adult NOD/scid gamma-null (NSG) mice were procured from the Jackson Laboratory, Bar Harbor, ME, USA (RRID:IMSR_JAX:001303). The mice were maintained and handled in accordance with approved protocols established by Geneva University animal facility.

### Reagents

Lys05 (Cat#SML2097), SAR405 (Cat#5.33063) and Ferric ammonium citrate (FAC, Cat#F5879) were from Sigma-Aldrich. Compound 9a was discovered and synthesized by J. Xu and Q. Gu.

### Culture of primary normal hematopoietic and leukemic cells

Viably frozen samples obtained from patients diagnosed with AML or normal CD34+ hematopoietic progenitor cells were thawed in 37°C water bath as fast as possible and placed immediately in 20% FCS Iscove’s Modified Dulbecco’s Medium (Gibco IMDM, Cat#21980-032, Thermo-Fischer Scientific) supplemented with 10μg/mL DNASE I (Sigma-Aldrich, Cat#11284932001). Next, cells were spun at 1,500 rpm for 5 min and re-suspended in complete medium (see below) at 2-5x10^6^ cells/mL. The thawed cells were subsequently cultured in complete medium: IMDM supplemented with BIT 9500 Serum Substitute (Cat#09500, StemCell Technologies) and BIT (bovine serum albumin 4 g/L, insulin 5 µg/mL, and transferrin 60 µg/mL, all from Sigma-Aldrich, Saint Louis, MO, USA). To facilitate optimal cell growth and maintenance, the supplemented medium, known as IMDM-BIT, was prepared with the addition of specific components. The following factors and reagents were incorporated into the IMDM-BIT medium: 50 ng/mL FLT3 ligand (Cat#300-19), 10 ng/mL IL-6 (Cat#200-06), 50 ng/mL stem cell factor (SCF, Cat#300-07), 25 ng/mL thrombopoietin (TPO, Cat#300-18), 10 ng/mL IL-3 (Cat#200-03), and 10 ng/mL granulocyte colony-stimulating factor (G-CSF, Cat#300-23) (all from Peprotech, Rock Hill, NJ, USA). Additionally, 50 µM β-mercaptoethanol (Sigma-Aldrich) was included in the culture medium.

### Patient-derived xenograft assays

To establish a humanized mouse model for studying AML, adult NOD/scid gamma-null (NSG) mice aged 6-8 weeks were utilized. The mice were treated with 20 mg/kg busulfan (Busilvex, Sigma-Aldrich) through intraperitoneal administration. Two days following busulfan treatment, viable primary human AML cells obtained from patients were injected into the tail vein at a range of 0.5-2x10^6^ cells. To evaluate the engraftment of human AML cells within the murine bone marrow hematopoietic environment, mice were monitored for varying durations, typically between 10 to 16 weeks. At the designated endpoint, the mice were humanely sacrificed, and the presence of viable human CD45^+^CD33^+^ cells was quantified using flow cytometry.

### Flow cytometry and cell sorting

To detect specific dyes and/or molecules using antibodies, flow cytometry analysis was performed using a Cytoflex flow cytometer (Beckman Coulter, Brea, CA, USA), following established protocols as previously described (*66*). For the identification and sorting of human AML cells, hCD33 and hCD45 staining were employed on an Astrios cell sorter (Beckman Coulter). Antibodies and dyes used are listed in the Supplementary Methods Section.

### In vivo cell kinetics assays

#### CFSE labeling

AML cells were subjected to a washing step using PBS. Subsequently, 1 µL of CFSE (CellTrace, Invitrogen, Carlsbad, CA, USA) stock solution was added to 1 mL of cell culture containing 0.5-5x10^6^ cells/mL. The cells were then incubated in the dark at 37°C for 30 minutes. Following this incubation, 50 mL of culture medium supplemented with 10% FBS was added to the cells, and they were further incubated for 5 minutes at room temperature. Subsequently, the cells were pelleted by centrifugation and resuspended in pre-warmed culture medium at 37°C.

#### BrdU labelling

When required, BrdU (Sigma-Aldrich) was administrated by intraperitoneal injection (100 mg/kg) twice a day. BrdU incorporation into human leukemic cells was measured using the APC BrdU Kit (BioLegends, San Diego, CA, USA) following the manufacturer’s instructions.

### Gene expression profiling

RNA quality was evaluated with a Bioanalyzer 2100 (using an Agilent RNA6000 nano chip kit, Santa Clara, USA), and 100 ng of total RNA was reverse transcribed using the GeneChip WT Plus Reagent Kit according to the manufacturer’s instructions (Affymetrix, Thermo-Fisher Scientific). Raw fluorescence intensities were normalized and analyzed as detailed in the Supplementary Methods Section.

### Single-Cell RNA sequencing

Single-Cell RNA sequencing was performed and analyzed as reported (*67*) and as detailed in the Supplementary Methods Section.

### Survival analysis

#### Analysis of AML cohorts

AML data from TCGA were obtained from cbioportal.org. Data from GSE14468 and BEAT AML were obtained from the GEO database. Briefly, raw counts were converted to TPM (transcripts per million) values following TMM (trimmed mean of M-values) adjustment in edgeR (version 3.32.1). Signature values were calculated by determining the mean expression of the signature genes. Survival analyses were performed using Cox regression from the survival package (version 3.2), and p-values were calculated using the log-rank test against the continuous signature value. Kaplan-Meier curves were plotted using a median cut-off.

#### Development of the QS35 risk score

We processed the genes upregulated within the q_clusters of leukemic cells (identified by scRNA-seq as enriched in Q_AML_UP and LSC-related signatures) with the glmnet package v4.1-3 (*68*) for searching correlations with survival in a training data set of 450 patients with AML (TCGA and GSE14468 cohorts). The validation cohort was identified within the BEAT AML database as patients with AML having received a comparable intensive therapy.

### Autophagic flux

Autophagic flux was assessed using the CYTO-ID® Autophagy Detection Kit (Enzo Life Science, Farmingdale, NY, USA) in accordance with the manufacturer’s instructions. Briefly, cells were initially labeled with suitable surface antibodies or Pyronin Y, and subsequently incubated at 37°C for 30 minutes with the Cyto-ID probe (1µl of Cyto-ID per 1ml of PBS). This incubation was performed either in the presence or absence of lys05 (5µM) to impede the autophagic flux. The cells were then washed with PBS, suspended in PBS containing a viability dye (DAPI or DRAQ7^+^), and subjected to flow cytometry analysis.

### Proteomics

Proteomics ESI-LC-MSMS assays were performed as reported (*67*) and as described in the Supplementary Methods Section. The mass spectrometry proteomics data have been deposited to the ProteomeXchange Consortium via the PRIDE partner repository with the dataset identifier PXD042796 (provisory access: Username; Password: R8EbaAfX).

### Vectors

We cloned the following oligonucleotides into the pLKO.1 mCherry constitutive vector (Dr Oskar Laur’s lab, Cat#128073; RRID:Addgene_128073) : control shRNA (CAACAAGATGAAGAGCACCAA), shNCOA4_1 (TCAGCAGCTCTACTCGTTATT) and shNCOA4_2 (TGAACAGGTGGACCTTATTTA).

### Statistics

Statistical analyses were conducted using Prism software 9.1.2 (GraphPad, San Diego, CA, USA). The differences between mean values obtained for the experimental groups were assessed using the two-tailed Student’s t-test with Welch’s correction or paired t-test when appropriate. For comparisons involving more than two groups, analysis of variance (ANOVA) was employed. Specific details, including the statistical tests used, the exact value of n, and the definition of measures, can be found in the figure legends. Typically, mean values and standard deviations were reported, and statistical significance was considered for p-values ≤0.05 and illustrated as follows: *p<0.05, **p<0.01, ***p<0.001.

## List of Supplementary Materials

Material and Methods

Fig. S1 to S6:

### Data files S1 and S2 (Excel files)

Data File S1. LSC signatures from literature (related to Figure 2). The list of genes defining the following

Data File S2. Gene sets defining the molecular signatures developed in this study (related to Figure 2).

## ACKNOWLEDGMENTS

We thank the Geneva flow cytometry, bioimaging, genomics (iGE3), proteomics, Reader Assay Development and Screening (READS) and zootechnie core facilities from Geneva University for technical support. We express our gratitude to Dr. Andrew A. Lane (Dana-Farber Cancer Institute in Boston, MA, USA) for engaging in valuable discussions on the manuscript.

## Author contributions

Conceptualization: C.L. and J.T. Methodology: C.L., P.A., S.M., P.T. and J.T.

Investigation: C.L., S.M., P.A., M.S., R.B., F.V., C.R., V.M.DM., G.Q*. and J.X*.

Visualization: C.L., P.A., S.M., P.T. and J.T.

Formal analysis: C.L., P.A., P.T., J.E.S. and J.T. Validation: C.L. and J.T.

Project administration: J.T. Supervision: C.L. and J.T.

Funding acquisition: C.L., J.E.S. and J.T. Writing – original draft preparation: J.T.

Writing – review and editing: C.L., P.T., J-E.S. and J.T.

*\* Discovered and synthesized compound **9a***.

## Data and materials availability

The raw and processed datasets from microarrays and scRNA-seq are available under accession number GEO: GSE235070 (provisory access using the reviewer token: kxshmqssljstbkl). Proteomic raw and processed data are accessible on to the ProteomeXchange Consortium via the PRIDE partner repository with the dataset identifier PXD042796 (provisory access: Username; Password: R8EbaAfX).

The original codes generated during this study can be made available upon reasonable request to the lead contact, Jerome Tamburini.

Any additional information required to reanalyze the data reported in this paper is available from the lead contact upon request.

## SUPPLEMENTARY MATERIAL AND METHODS

### Animals

Upon arrival, the mice had an average weight of 20-30g and were housed in cages with littermates, accommodating 5-6 mice per cage. Standard conditions, including temperature, humidity, and lighting, were maintained throughout the study. The mice were provided with a standard chow diet and had ad libitum access to food and water. To ensure unbiased and controlled experimental conditions, the assignment of mice to respective experimental groups was randomized. Littermate controls were included for comparative analysis. Throughout the study, all animal handling, treatment administration, and euthanasia procedures were performed in strict accordance with the approved protocols established by the Geneva health department.

### Cell Lines

Cells were cultured in a 37°C incubator with a 5% CO2 atmosphere. The HEK293T/17 cell line was utilized for lentiviral production. Cells were cultured in Gibco Dulbecco’s modified Eagle medium (DMEM) Glutamax (Cat#61965, Thermo-Fischer Scientific, Waltham, MA, USA) supplemented with 10% fetal calf serum (FCS, Eurobio Scientific, Cat#CVFSVF00-01). The murine stromal cell line, MS5, was employed in co-culture assays with human leukemic cells. MS5 cells were cultured in DMEM Glutamax supplemented with 10% FCS. The MOLM-14 human AML cell line is short tandem repeat (STR; PCR-single-locus-technology, Promega PowerPlex21 PCR Kit, Eurofins Genomics) profiled periodically, and verified Mycoplasma negative yearly.

### Flow Cytometry Antibodies and Dyes

FITC Mouse Anti-Human CD45 Cat# 555482 RRID:AB_395874; APC Mouse Anti-Human CD45 Cat# 555485 RRID:AB_398600; CD33 PerCP-Cy™5.5 Cat# 333146 RRID:AB_2868647; PE Mouse Anti-Human CD33 Cat# 555450 RRID:AB_395843; PE Rat Anti-Mouse CD45 Cat# 553081 RRID:AB_394611; PE Mouse Anti-Mouse CD45.1 Cat# 553776 RRID:AB_395044; FITC Mouse Anti-Human CD33 Cat# 555626 RRID:AB_395992; AF647 Mouse anti-Ki-67 Cat# 561126 RRID:AB_10611874; FITC Mouse Anti-Ki-67 Cat# 556026 RRID:AB_396302; APC Mouse Anti-Human CD71 Cat# 551374 RRID:AB_398500; BV421 Mouse Anti-Human CD34 Cat# 562577 RRID:AB_2687922; PE Mouse Anti-Human CD34 Cat# 555822 RRID:AB_396151; APC Mouse Anti-Human CD38 Cat# 555462 RRID:AB_398599; PE-Cy7 Mouse Anti-Human CD38 Cat# 335790 RRID:AB_399969; FITC Mouse Anti-BrdU Cat# 556028 RRID:AB_396304 from BD Biosciences, APC anti-BrdU Cat# 364113 RRID:AB_2814314 and APC anti-human Ki-67 Cat# 350513 RRID:AB_10959326 from BioLegend. Dyes used were: DAPI Axonlab Cat#A4099.0, CellTrace CFSE Cell Proliferation Kit (Life Techniologies, Cat#C34570), Pyronin Y (Sigma Aldrich, Cat#P9172), Hoechst 33342 (Thermo Fisher Scientific, Cat#62249), 5-Bromo-2′-deoxyuridine (Sigma Aldrich, Cat#B5002), Propidium iodine (BD Biosciences, Cat#556463, RRID:AB_2869075), Cyto-ID (Enzo Life Sciences, Cat#ENZ-51031), BioTracker Labile Fe^2+^ (Merck Millipore, Cat#SCT037), Transferrin From Human Serum, Alexa Fluor™ 488 Conjugate (Thermo Fisher Scientific, Cat#T13342), Annexin V (BD Biosciences, Cat#556419, RRID:AB_2665412) and DRAQ7^+^ (Thermo Fisher Scientific, Cat#D15106). The following commercially available kits were used: BD Cytofix/Cytoperm Kit (BD Biosciences, Cat#554714, RRID:AB_2869008), BD Cytoperm Permeabilization Buffer Plus (BD Biosciences, Cat#561651), CellROX Deep Red (Thermo Fisher Scientific, Cat#C10422), MitoTracker Green (Thermo Fisher Scientific, Cat#M7514), MitoTracker Deep Red (Thermo Fisher Scientific, Cat#M22426), LysoTracker Deep Red (Thermo Fisher Scientific, Cat#L12492), MitoSox Red (Thermo Fisher Scientific, Cat#M36008) and MitoSox Green (Thermo Fisher Scientific, Cat#M36006).

### Cell cycle analysis

#### Ki67/DAPI staining

AML cells were initially fixed in 4% paraformaldehyde (Sigma-Aldrich) for 15 minutes at room temperature. Following fixation, the cells were washed twice with PBS and subsequently permeabilized using cold methanol for 2 hours on ice. After an additional two washes with PBS, the cells were stained with an anti-Ki67 antibody (Becton Dickinson, BD, Franklin Lakes, NJ, USA) for 1 hour at room temperature. Subsequently, the cells were washed twice with PBS, resuspended in PBS containing DAPI (Thermo-Fischer Scientific), and subjected to analysis using a Cytoflex flow cytometer (Beckman Coulter).

#### Pyronin Y/Hoechst Staining

AML cells were washed with PBS and then incubated at 37°C for 45 minutes with 20 μg/mL Hoechst 33342 (Invitrogen) in Hanks balanced salt solution (HBSS) supplemented with 10% FCS, 20 mM Hepes, and 50 µg/mL verapamil at pH 7.5. Next, 1 µg/mL Pyronin Y (Sigma-Aldrich) was added, and the cells were incubated for an additional 15 minutes at 37°C. The cells were promptly processed for analysis using a Cytoflex cytometer or sorted using an Astrios cytometer.

### Gene expression profiling

#### Gene chip hybridization

RNA was extracted using a RNeasy Mini Kit (Cat#74134, Qiagen, Redwood City, CA, USA) and quality was evaluated with a Bioanalyzer 2100 (using Agilent RNA6000 nano chip kit), and 100 ng of total RNA was reverse transcribed using the GeneChip® WT Plus Reagent Kit according to the manufacturer’s instructions (Affymetrix, Thermo Fischer Scientific). Briefly, cDNA were hybridized to GeneChip® Clariom S Human (Affymetrix) at 45°C for 17 hours, then washed on the fluidic station FS450 (Affymetrix), and scanned using the GCS3000 7G (Thermo Fischer Scientific). Scanned images were then analyzed with Expression Console software (Affymetrix) to obtain raw data (.cel files) and metrics for quality controls.

#### Data processing

The raw fluorescence intensity values were subjected to a series of normalization steps, including background correction, quantile normalization, and log2 transformation, utilizing the Transcriptome Analysis Console (TAC) v4.0.2 (Thermo-Fischer Scientific). To discern genes exhibiting differential expression, an analysis of variance (ANOVA) approach, accompanied by empirical Bayesian (eBayes) correction, was employed. The cutoff criteria for fold-changes (FC) between the comparison groups are provided in the relevant sections.

#### Molecular signatures

The statistical analyses were conducted using the R environment for statistics (v.4.0.2). The gene expression data from the remaining samples were subjected to analysis using the limma package (v.3.46.0). The analysis design incorporated patient-specific effects and differentiated between quiescent and proliferative cell states. Genes exhibiting a log2-fold change ≥ │1│ and an adjusted p-value (FDR) below 0.1 were selected for subsequent analysis and application to additional datasets. To mitigate technical bias, the top gene list excluded histone genes, as they were frequently absent in TCGA RNAseq data. Signature gene sets were established based on genes upregulated in quiescent (Q_AML_UP) and cycling (C_AML_UP) cells, respectively.

#### Gene set enrichment analysis

To identify differentially expressed genes, we applied a fold change filter, setting it to at least a ≥1.5 or ≤-1.5 fold difference in expression between groups with an FDR threshold of <0.05. The resulting data were then analyzed using Gene Set Enrichment Analysis (GSEA) from the Broad Institute to investigate the dysregulation of biological pathways. Enrichment rates with a P-value of <0.05 and FDR ≤0.1 were considered significant.

#### Pathway analysis

We conducted pathway analysis by utilizing a ranked gene list derived from differential gene expression analysis between quiescent and cycling leukemic cell fractions. GSEA was carried out with default parameters, employing 1000 permutations. For this analysis, we utilized the pathway database c5.go.v2023 from GSEA. The resulting data were visually represented in Cytoscape 3.9.1 using EnrichmentMap. To facilitate comprehension, clusters were generated and annotated using AutoAnnotate. Furthermore, manual adjustments were made to cluster labels to improve clarity.

### Single-Cell RNA sequencing

After lysing red blood cells with EL buffer (Qiagen, Cat#79217), a total of 10^7^ cells isolated from mouse bone marrow of patient-derived xenografts (PDXs) were subjected to staining. Fixable Viability Stain 510 and hCD45-APC and hCD33-PE antibodies were used for staining, following the manufacturer’s instructions. Viable cells positive for hCD45 and hCD33 were isolated using an Astrios cell sorter and suspended in PBS supplemented with 0.04% Ultrapure BSA (Invitrogen, Cat# AM2616) at a concentration of 1500 cells/µL. Subsequently, up to 10^5^ cells per sample were encapsulated using the 10X Chromium controller (10X Genomics) according to the Chromium Next GEM Single Cell 3ʹ User Guide v3.1 (10X Genomics). In brief, the cells were enclosed within oil droplets along with barcoded Gel Beads and reagents for cDNA synthesis from mRNA. The resulting cDNAs underwent amplification, fragmentation, and ligation with Illumina adapters. A final index PCR was performed, and the libraries’ quality was assessed using the Tapestation 2200 system (Agilent, Santa Clara, CA. USA). The libraries were then frozen prior to quantitation and sequencing. Barcoding was implemented at three levels: cell barcodes were assigned to each sequence read to identify its corresponding cell of origin, unique molecular identifiers (UMIs) located upstream of poly(d)T primers allowed tagging of each original molecule to mitigate amplification bias, and an index enabled pooling of different samples. Quantification of the single-cell RNA sequencing (scRNA-seq) libraries was performed using HSD1000 reagents on the TapeStation 2200 system (Agilent), and the libraries were pooled at equimolar concentrations within a size range of 200 to 700 base pairs. The pooled libraries were then sequenced at a concentration of 1.8pM with 1% phiX on a NextSeq 500 sequencer (Illumina, San Diego, CA, USA) to achieve a sequencing depth of 40,000-50,000 reads per cell. The sequencing configuration included paired-end reads with 28 cycles for the first read, 91 cycles for the second read, and 8 cycles for a single index.

#### Data analysis

Single-cell RNA sequencing (scRNA-seq) data output files in bcl2 format were converted to FASTQ format using Cell Ranger v.6.1.1.1 (ref). The resulting count matrices from Cell Ranger were imported into the R programming environment v4.1.2 using the Seurat package v4.1.0. To ensure data quality, cells with a low number of unique genes (<7500 unique genes) and a high proportion of mitochondrial RNA (>20%) were removed. Gene counts were then normalized using a regularized negative binomial regression method, employing the SCT normalization workflow from the Seurat package (ref). The integration process utilized 30 principal components and 3000 genes. Subsequently, dimensional reduction was performed, summarizing the 30 principal components using UMAP dimensionality reduction. Cell clusters were identified through Louvain clustering within the Seurat workflow, with the resolution parameter set to 0.25. Signature values on the SCT-transformed data were computed using the AddModuleScore method from Seurat. Furthermore, the nclust package was utilized to perform gene and cell clustering, employing Pearson’s correlation similarity and average linkage. To facilitate heatmap visualization, the normalized single-cell expression data underwent mean centering. A distinct cell cluster exhibiting a quiescent phenotype was identified by considering the high expression of Q_AML_UP genes, low expression of C_AML_UP genes, and reference to previously published signatures (**Table S1**). To facilitate further analyses, genes specific to the identified quiescent clusters (referred to as q_clusters) were extracted. These q_cluster genes were characterized by weak expression in other cells but demonstrated a strong correlation within the quiescent population.

### Proteomics

#### Sample preparation

Cell pellets were collected and resuspended in 100 μl of 0.1 % RapiGest Surfactant (Waters Corporation, Milford, MA, USA) in 0.1M triethylammonium bicarbonate (TEAB). Samples were heated for 5 min at 95°C, followed by lysis using sonication (6 x 30 sec.) at 70% amplitude and 0.5 pulse, with 30 sec. on ice between each cycle of sonication. Samples were centrifuged and the remaining pellet was solubilized in 100 μl of 0.1 % RapiGest with 2 μL of 0.1M MgCl2 and 0.2 μL of benzonase for DNA digestion. The protein lysates were pooled together, and protein concentration was measured by Bradford assay. A total of 25 μg of each sample was digested with trypsin, as follows: sample volume was adjusted to 100 μl with 0.1M TEAB to obtain a final concentration of RapiGest 0.1%, followed by reduction with 2 μl of Dithioerythritol (DTE) 50 mM in distilled water at 60°C for 1h. Alkylation was then performed by adding 2 μL of iodoacetamide (400 mM in distilled water) for 1 hour at room temperature in the dark. Overnight digestion was performed at 37 °C with 5 μL of freshly prepared trypsin (Promega, Madison, WI, USA) at 0.1 μg/μl in TEAB 0.1M. To remove RapiGest, samples were acidified with TFA, heated at 37°C for 45 min, and centrifuged. Supernatants were desalted with a C18 microspin column (Harvard Apparatus, Holliston, MA, USA) according to manufacturer’s instructions, dried completely under speed-vacuum, and stored at -20°C.

#### ESI-LC-MSMS

Samples were prepared by diluting them in 25 μL of loading buffer (5% CH3CN, 0.1% FA) and adding 1.5 μL of Biognosys iRT peptides to each sample. Two microliters of the prepared sample were injected onto a column for LC-ESI-MS/MS analysis. The analysis was performed on an Orbitrap Fusion Lumos Tribrid mass spectrometer (Thermo Fisher Scientific) equipped with an Easy nLC1200 liquid chromatography system (Thermo Fisher Scientific). The peptides were trapped on a Acclaim pepmap100, C18, 3μm, 75μm x 20mm nano trap-column (Thermo Fisher Scientific) and separated on a 75 μm x 500 mm, C18 ReproSil-Pur (Dr. Maisch GmBH, Germany), 1.9 μm, 100 Å, home-made column. The separation was carried out over 135 min using a gradient of H2O/FA 99.9%/0.1% (solvent A) and CH3CN/H2O/FA 80.0%/19.9%/0.1% (solvent B). Data-Independent Acquisition (DIA) was used with MS1 full scan at a resolution of 60,000 (FWHM), followed by 30 DIA MS2 scans with variable windows. MS1 was performed in the Orbitrap with an AGC target of 1 x 10^6^, a maximum injection time of 50 ms, and a scan range from 400 to 1240 m/z. DIA MS2 was performed in the Orbitrap using higher-energy collisional dissociation (HCD) at 30%. The isolation window was set to 28 m/z with an AGC target of 1 x 10^6^ and a maximum injection time of 54 ms.

#### Data analysis

DIA raw files were loaded into Spectronaut v.14 (Biognosys, Schlieren, Switzerland) and analyzed by directDIA using default settings. Briefly, data were searched against Human reference proteome fasta database (Uniprot, release 2020_09, 55471 entries). Trypsin was selected as the enzyme, with one potential missed cleavage. Variable amino acid modifications were oxidized methionine and deaminated (NQ). Fixed amino acid modification was carbamidomethyl cysteine. Both peptide precursor and protein FDR were controlled at 1% (Q value < 0.01). Single Hit Proteins were excluded. For quantitation, Top 3 precursor area per peptides were used, “only protein group specific” was selected as proteotypicity filter and normalization was set to “global normalization”. The quantitative analysis was performed with MapDIA tool, using the precursor quantities extracted from Spectronaut output. No further normalization was applied. The following parameters were used: min peptides = 2, max peptides = 10, min correl = -1, Min_DE = 0.01, max_DE = 0.99, and experimental_design = replicate design. Proteins were considered to have significantly changed in abundance with an FDR ≤ 0.05 and an absolute fold change FC≥ |1.5| (log2FC ≥ |0.58|). FDR q-value was calculated using the two-stage step-up method of Benjamini, Krieger and Yekutieli (Graphpad Prism). Proteins were excluded when more than one replicate was impaired per condition. If only one replicate was impaired, it was excluded and the protein conserved. The mass spectrometry proteomics data have been deposited to the ProteomeXchange Consortium via the PRIDE partner repository with the dataset identifier PXD042796 (provisory access: Username; Password: R8EbaAfX).

### Clonogenic assays

#### L-CFU assays

L-CFU assays were performed as previously described (*69*). Briefly, AML cells were seeded at 10^5^/mL in H4230 medium (Cat#04230, StemCell Technologies, Vancouver, Canada) supplemented with 10% of the following medium: IMDM, 50 ng/mL FLT3L, 10 ng/mL IL-6, 50 ng/mL SCF, 25 ng/mL TPO, 10 ng/mL IL-3, and 10 ng/mL G-CSF. At day 7, L-CFU (colony of > 10 cells) were scored under an inverted microscope.

#### Normal hematopoietic progenitor clonogenic assays

Normal CD34^+^ hematopoietic cells (Cat#70002.2, StemCell Technologies) were seeded at 10^4^/ml in MethoCult H4034 Optimum medium (Cat#04064, StemCell Technologies). The erythroid burst-forming units (BFU-E) and granulocyte-macrophage colony-forming units (CFU-GM) were counted under an inverted microscope at day 7 and 14.

#### LTC-IC assays

Leukemic cells were cultured in complete medium as described above and were subjected to shRNA transduction or treatment with various compounds as indicated in the corresponding sections. For the preparation of LTC-IC assays, poly-L-lysine-coated 96-well plates were seeded with 5x10^3^ irradiated MS-5 cells in a final volume of 100 µL per well. Subsequently, leukemic cells were seeded at different concentrations in Myelocult H5100 (Cat#05150, StemCell Technologies) supplemented with recombinant human IL-3, G-CSF, and TP0 cytokines (20 ng/mL each). The cells were cultured for a period of 5 weeks with medium replaced twice a week. After 5 weeks, the cells were harvested, and the contents of each well were seeded in 500 µL methylcellulose in 24-well culture plates. Colonies consisting of at least 10 cells were counted under an inverted microscope after 10-14 days. A well was considered positive if it gave rise to at least one colony. The frequency of LTC-IC was determined using the ELDA software.

### Lentiviral production

We cloned the following oligonucleotides into the pLKO.1 mCherry constitutive vector (Dr Oskar Laur’s lab, Cat#128073; RRID:Addgene_128073) : control shRNA (CAACAAGATGAAGAGCACCAA), shNCOA4_1 (TCAGCAGCTCTACTCGTTATT) and shNCOA4_2 (TGAACAGGTGGACCTTATTTA). HEK293T/17 cells were transfected with the packaging plasmids pMD2.G (Dr Didier Trono’s lab, Cat#12259; RRID:Addgene_12259) and psPAX2 (Dr Didier Trono’s lab, Cat#12260; RRID:Addgene_12260) and the pLKO.1 mCherry constitutive vectors expressing shRNAs, all amplified in NEB® Stable Competent E. coli (NEB, Cat#C3040H) using Lipofectamine 2000 Transfection Reagent. After 24 hours, cell culture medium was changed and optimized Minimal Essential Medium (Gibco opti-MEM, Cat#31985-047, Thermo-Fischer Scientific) was added, allowing the cells to produce lentiviral particles. The culture supernatants from HEK293T/17 cells containing lentiviral particles were collected 72 hours after transfection, filtered, concentrated using PEG-it Virus Precipitation Solution (PEG-it Virus Precipitation Solution, System Biosciences, Palo alto, CA, USA, Cat#LV810A), following the manufacturer’s instructions, and stored at -80 °C.

### Lentiviral infection

Twelve-well plates were coated with 500 µL of retronectin (20 µg/mL in PBS, Takara Bio, Cat#T100A) and incubated overnight at 4°C. Subsequently, the plates were blocked with 1 mL of 2% BSA PBS for 30 minutes at room temperature and washed once with PBS. Next, 600 µL of concentrated lentiviral supernatant was added to each well, and the plates were centrifuged at 4000 rpm for 3 hours at 4°C to enhance virus adhesion to retronectin. Following centrifugation, 0.4 mL of the viral supernatant was replaced with 1 mL of pre-warmed culture medium containing 1-5×10^6^ cells. The plates were then centrifuged at 1500 rpm for 10 minutes at room temperature and incubated at 37°C for 24 hours.

### Bioenergetics analysis experiments

Oxygen consumption was measured using a Cell Mito Stress Test kit (Agilent Technologies) on a Seahorse XF24 extracellular flux analyzer. Seahorse XFp microplate wells were coated with 25 µl of Cell-Tak solution at a concentration of 22.4 µg ml−1 and kept at 4 °C overnight. Then, 5 × 10^5^ cells per well were adhered to Seahorse microplates and rested for 1 h without CO_2_ in XF base minimal DMEM containing 11 mM glucose, 1 mM pyruvate and 2 mM glutamine. Then, wells were successively injected with oligomycin (inhibitor of ATP synthase), carbonilcyanide p-triflouromethoxyphenylhydrazone (FCCP, a decoupling agent that disrupts the mitochondrial membrane potential) and combination of rotenone and Tntimycin A (Rot/AA that inhibit mitochondrial complexes I and III, respectively).

### Softwares

GraphPad Prism http://www.graphpad.com/, RRID:SCR_002798; FlowJo https://www.flowjo.com/solutions/flowjo RRID:SCR_008520; R language for statistics http://cran.r-project.org RRID:SCR_001905; Seurat R package https://satijalab.org/seurat/ RRID:SCR_016341; GLMnet package https://glmnet.stanford.edu/articles/glmnet.html RRID:SCR_015505; Nclust package https://gitlab.com/pwirapati/nclust; Transcriptome Analysis Console (TAC); Thermo-Fischer Scientific RRID:SCR_016519; ELDA https://bioinf.wehi.edu.au/software/elda/ RRID:SCR_018933; EnrichmentMap version 3.3.6 https://apps.cytoscape.org/apps/enrichmentmap RRID:SCR_016052; Cytoscape 3.8.2 https://cytoscape.org/ RRID:SCR_003032 and Biorender https://www.biorender.com/ RRID:SCR_018361.

---

**Figure S1.**
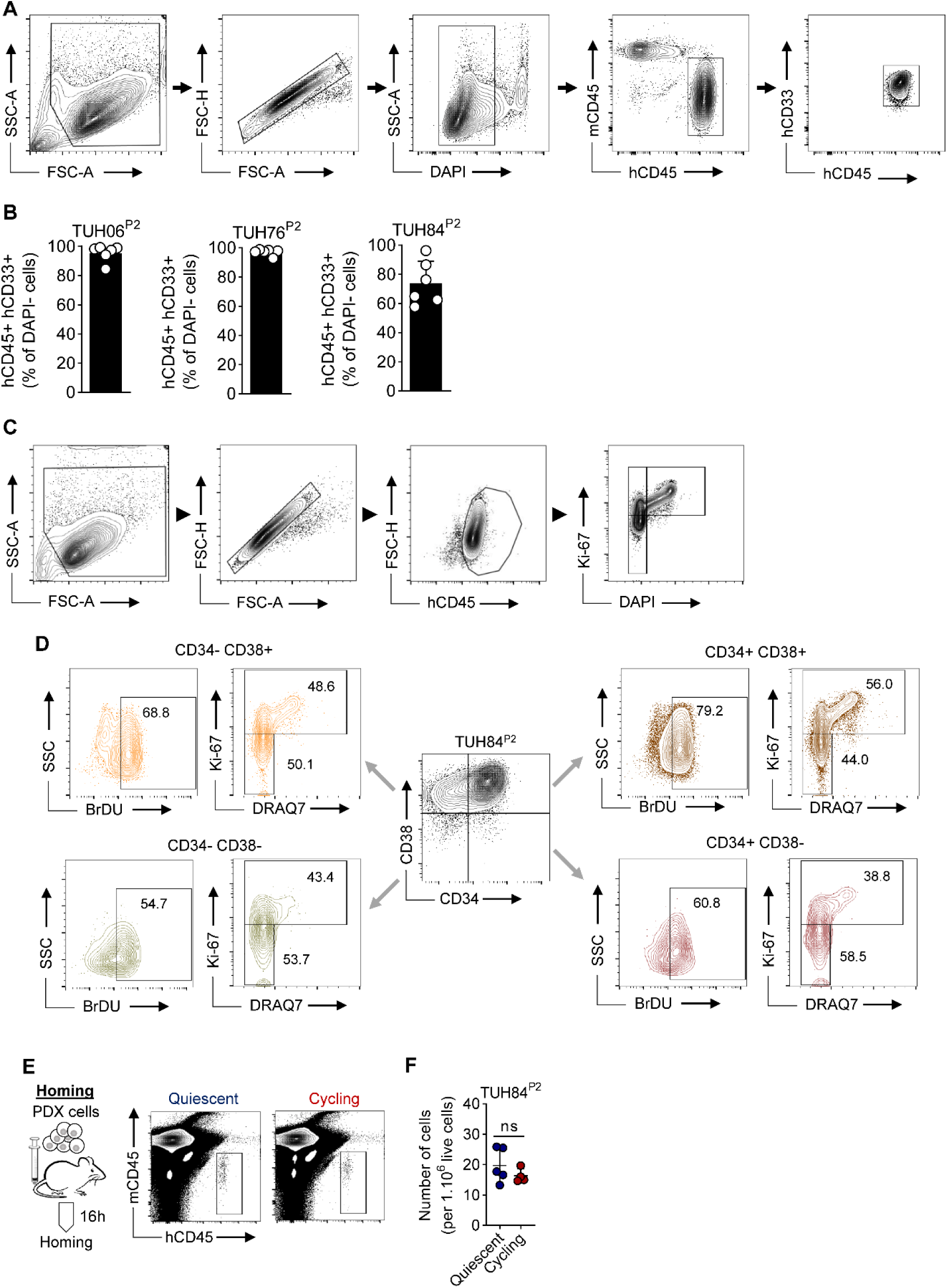
Flow Cytometry Gating Strategies on Human AML cells in PDX Models. **A.** Gating strategy for human AML cells identification within murine bone marrow samples from PDX assays. Side scatter area (SSC-A), forward scatter area (FSC-A), forward scatter height (FSC-H), anti-CD45 antibody with human reactivity (hCD45), anti-CD45 antibody with reactivity against mice (mCD45), and anti-CD33 antibody with reactivity against human (hCD33) were utilized in murine bone marrow samples from PDX assays. **B.** Quantification of human CD33^+^CD45^+^ AML cells among viable DAPI^-^ cells in murine bone marrow samples (n=2 different PDX assays, n=6 mice per assay). **C.** Gating strategy for cell cycle analysis using Ki67 and DAPI staining in human hCD45^+^ leukemic cells. **D.** Representative contour plots panels illustrating the gating strategy for Ki67/DRAQ7^+^ or BrdU staining among the four subpopulations defined by CD34 and CD38 staining in human AML cells from PDX assays. **E-F.** Cycling or quiescent populations were sorted using pyronin Y and Hoechst staining and subsequently transplanted separately into NSG mice. Mice were sacrificed after a 16-hour homing period, and bone marrow was harvested to assess human cell engraftment. **E.** Representative contour plots showing hCD45 versus mCD45 staining in the quiescent and cycling groups. **F.** Number of human leukemic cells (per 10^6^ live cells) among murine bone marrow cells. Error bars indicate standard deviations. "ns" denotes no significance.

**Figure S2.**
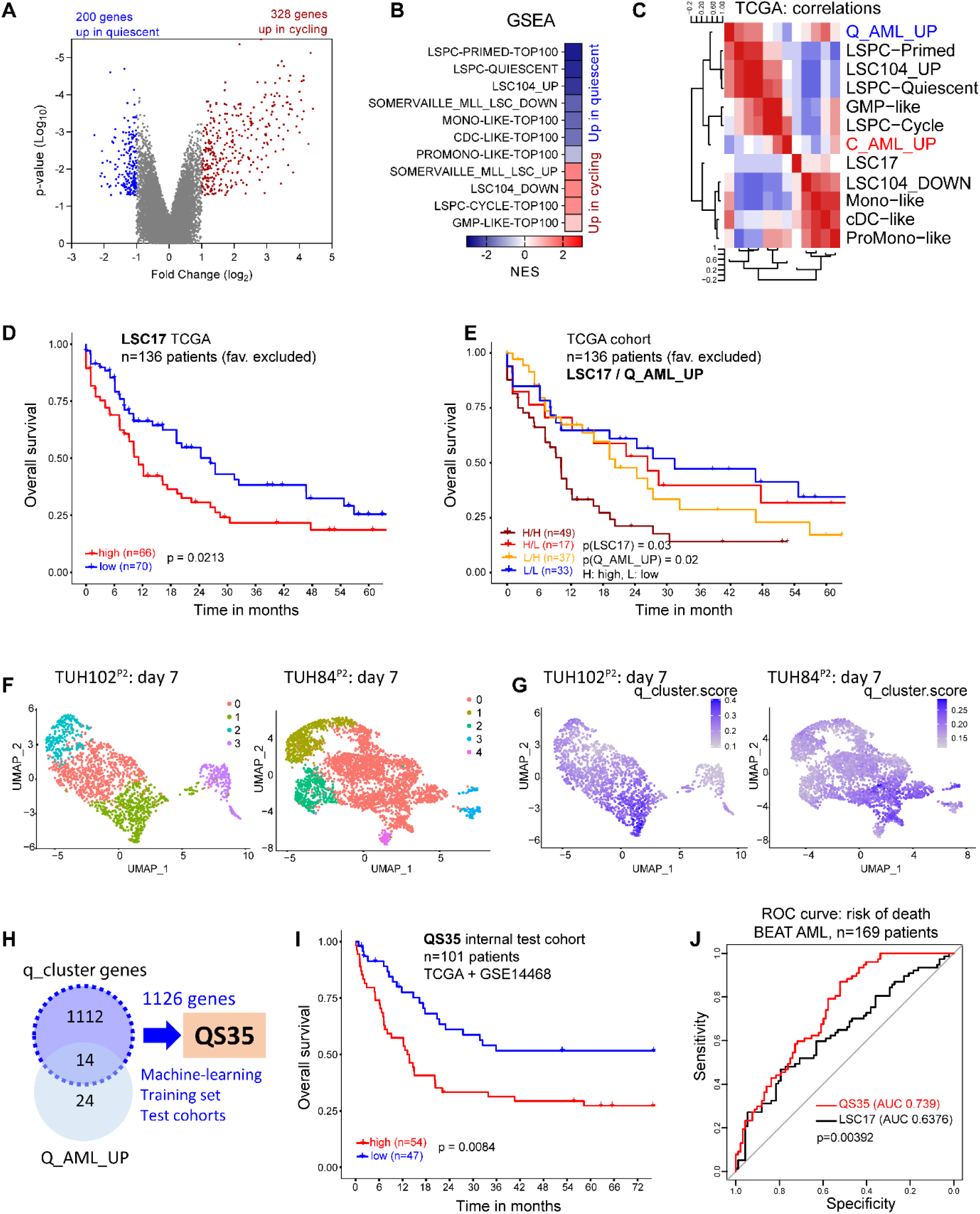
Molecular Attributes of Leukemia Stem Cells are Found in the Quiescent Subset of AML Cells. **A-B.** Sorting of AML cells into cycling (Pyronin Y^+^) and quiescent (Pyronin Y^-^) populations to perform differential gene expression (DGE) analysis and transcriptomic profiling in four distinct patient-derived xenografts (PDX). **A.** Volcano plot illustrating differential gene expression between cycling and quiescent transcriptomes. Genes upregulated in cycling or quiescent subsets (log2 fold-change (FC) ≥ |0.585|, p-value<0.05) are highlighted in red or blue, respectively, with the number of genes indicated. **B.** Enrichment analysis conducted for several leukemic stem cell-related signatures in cycling versus quiescent populations, with the normalized enrichment score (NES) displayed using a color scale. **C.** Correlation heatmap for various gene sets including the Q_AML_UP signature in the TCGA cohort. **D-E.** The TCGA data set was analyzed following the exclusion of patients classified under the favorable category (n=136 patients). **D.** Survival curves of the LSC17 signature within the TCGA cohort. p-values are provided when considering the signature as continuous variable, or with a median cut-off (the later in brackets). **E.** Survival curves within the TCGA cohort divided into four groups of patients based on high (H) or low (L) expression of the Q_AML_UP and LSC17 signatures. The number of patients within these subgroups is indicated. P-values are provided when considering the signature as continuous variable, or with a median cut-off (the later in brackets). **F.** Clusters derived with the Louvain Algorithm in Seurat are depicted using distinct colors. **G.** Uniform manifold approximation and projection (UMAP) plot for enrichment in the q_cluster score in the samples TUH102^P2^ and TUH84^P2^ generated using Seurat. **H.** Schematic representation of the development of the QS35 signature from single-cell AML transcriptomes. **I.** QS35 signature in the internal validation cohort constituted of 101 patients from both TCGA and GSE14468 cohorts. **J.** The Receiver Operating Characteristic (ROC) curves were plotted to evaluate the predictive performance of the QS35 and LSC17 signatures for assessing the risk of mortality within the validation cohort derived from BEAT AML. AUC: area under the curve. The p-values determined using the median cut-off are reported.

**Figure S3.**
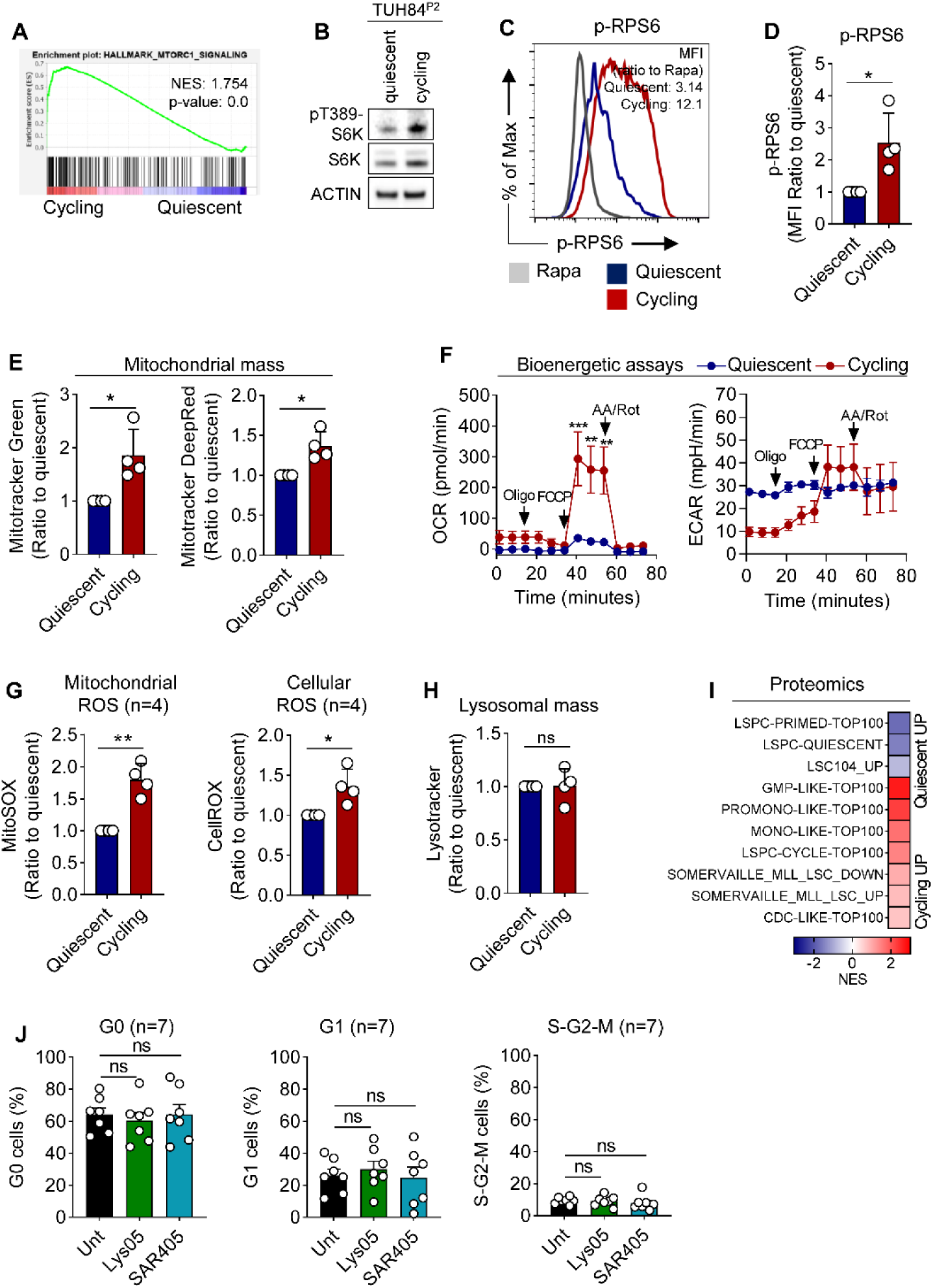
Characterization of Metabolic Pathways Activated in Cycling Compared to Quiescent Leukemic Cells. **A.** Gene set enrichment analysis (GSEA) for mTORC1 signaling comparing quiescent and cycling AML cells. Normalized enrichment score (NES) and p-values are provided. **B.** Quiescent and cycling PDX AML cells were sorted based on pyronin Y / Hoechst staining. Western blot analysis was performed using antibodies against anti-phospho-S6K (Thr389), -S6K and actin. **C-D.** AML cells were treated with either vehicle or 10 nM rapamycin for 24 hours to modulate mTORC1 activity. The phosphorylation level of the downstream target ribosomal protein S6 (RPS6) was assessed using flow cytometry. **C.** Representative univariate plots of phospho-RPS6 in quiescent, cycling, or rapamycin-treated AML cells were generated. **D.** Quantification of phospho-RPS6 was performed by calculating the ratio of each condition to the signal obtained in quiescent cells (n=4). **E.** Mitochondrial mass in quiescent and cycling fractions was measured using MitoTracker Green and DeepRed dyes (n=4). **F.** Bioenergetic assays were conducted to measure the oxygen consumption rate (OCR) and extracellular acidification rate (ECAR) in mpH/min in sorted quiescent or cycling AML cells. Oligomycin (O, mitochondrial ATP synthase inhibitor), Carbonyl cyanide-4 (trifluoromethoxy) phenylhydrazone (FCCP, mitochondrial uncoupling agent), and rotenone and antimycin (R, complex I and complex III inhibitors, respectively) were used as specific inhibitors of mitochondrial respiratory chain during the assays. **G.** Mitochondrial and total reactive oxygen species (ROS) production in quiescent and cycling fractions was measured using MitoSOX and CellROX staining, respectively (n=4). **H.** The lysosomal mass in cycling or quiescent fractions was assessed using the LysoTracker dye (n=4). **I.** Enrichment analysis was performed to identify differentially expressed proteins between the quiescent and cycling fractions. The normalized enrichment score (NES) was calculated and represented using a color scale. **J.** AML cells were incubated with vehicle, or 1 μM Lys05 or 5 μM SAR405 for 48 h. Cell cycle distribution between G_0_, G_1_ and S-G_2_-M subpopulations was assessed using Ki67 and DAPI staining (n=7). Error bars indicate standard deviations. "ns" denotes no significance, **p<0.01, ***p<0.001.

**Figure S4.**
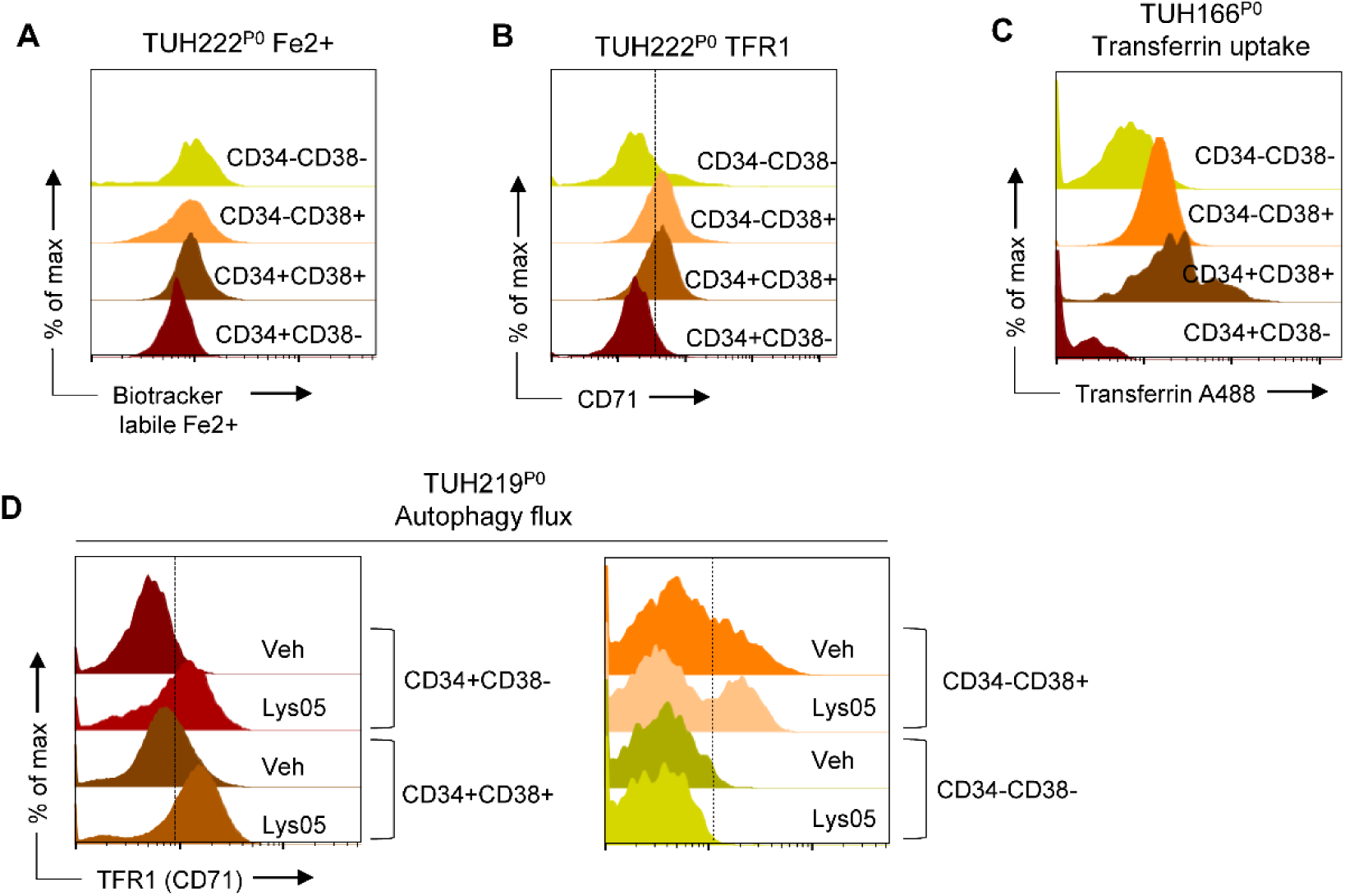
Low Iron Metabolism is Linked to Quiescent Leukemic Stem Cells in Primary AML Cells. **A-C.** Quantitative analysis of various parameters within each of the four leukemic cell subpopulations defined by CD34/CD38 markers in primary AML cells. Representative flow cytometry histograms are shown. **A.** Intracellular Fe^2+^ quantification using the BioTracker Labile Fe^2+^ dye. **B.** TFR1 expression quantification at the cell surface. **C.** Measurement of transferrin uptake using AF488-labelled transferrin. **D.** Leukemic cells were incubated with vehicle or 2 μM Lys05 for 12 h, and the four leukemic cell subpopulations defined by CD34/CD38 markers were analyzed for Cyto-ID staining.

**Figure S5.**
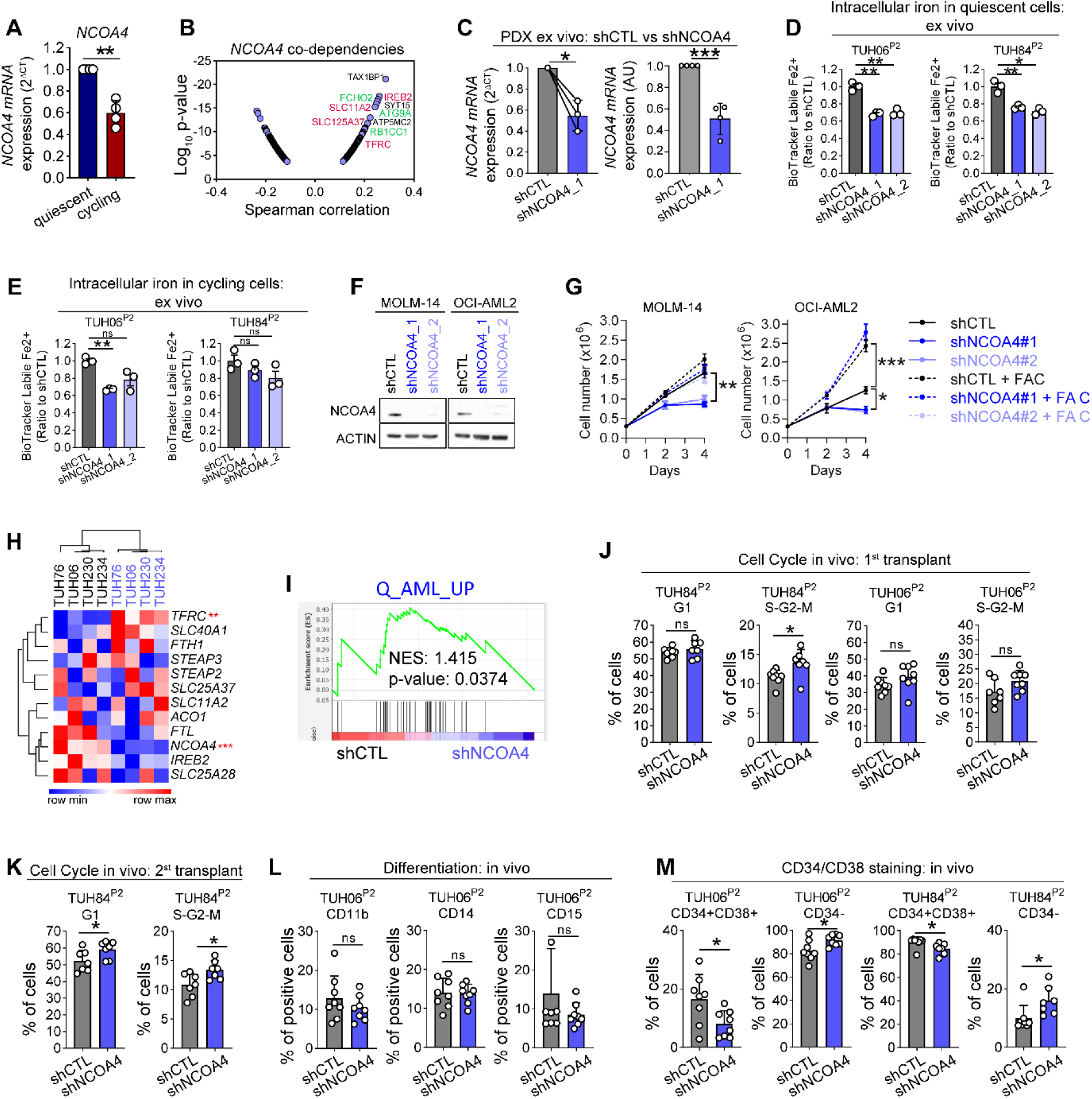
Multi-Dimensional Readouts Unveiling NCOA4 Depletion Effects in Human AML Cells. **A.** Quantification of *NCOA4* expression in sorted quiescent or cycling leukemic cells by quantitative PCR (n=4). **B.** Analysis of NCOA4 co-dependencies in cancer using the DepMap online tool. Genes relevant to iron metabolism (in red) or autophagy (in green) are highlighted. **C-M.** AML cells were transduced ex vivo with lentiviral vectors expressing mCherry-tagged NCOA4-targeting or control shRNAs. **C.** Quantification of *NCOA4* expression by quantitative PCR (left panel, n=3) and microarrays (right panel, n=4). **D-E.** Quantification of the intracellular labile iron pool using the BioTracker Labile Fe^2+^ dye in the quiescent (**D**) or cycling (**E**) subpopulations defined by Ki67 and DAPI staining ex vivo (n=3 replicates in two different PDXs). **F-G.** MOLM-14 and OCI-AML2 cell lines were transduced with control or anti-NCOA4 shRNAs. **F.** Western blots using anti-NCOA4 and anti-actin antibodies. **G.** Trypan blue exclusion assays during four days to measure the number of viable cells after incubation without or with 10 mg/mL FAC. **H-I.** Differential gene expression between NCOA4-depleted or control leukemic cells. **H.** Unsupervised clustering analysis for selected iron metabolism-related genes. Significant differences are highlighted. **I.** Enrichment in the Q_AML_UP signature in control versus NCOA4-depleted cells. **J-M.** AML cells transduced with control of anti-NCOA4 shRNAs were sorted based on Pyronin-Y/Hoechst staining, and the quiescent fraction was transplanted to immunodeficient NSG mice (n=7 mice per group in two different PDX assays). After 10-12 weeks in vivo, transplantation to secondary recipient mice were done (n=8 mice per group in one PDX assay). **J-K.** Percentage of human AML cells in G_1_ or S-G_2_-M phase of the cell cycle after the first (**J**) and the second transplantation (**K**). **L.** Cell membrane positivity for CD11b, CD14 and CD15 differentiation markers. **M.** The proportion of CD34^+^CD38^+^ and of CD34^-^ cells is provided. Error bars indicate standard deviations. "ns" denotes no significance, *p<0.05, **p<0.01.

**Figure S6.**
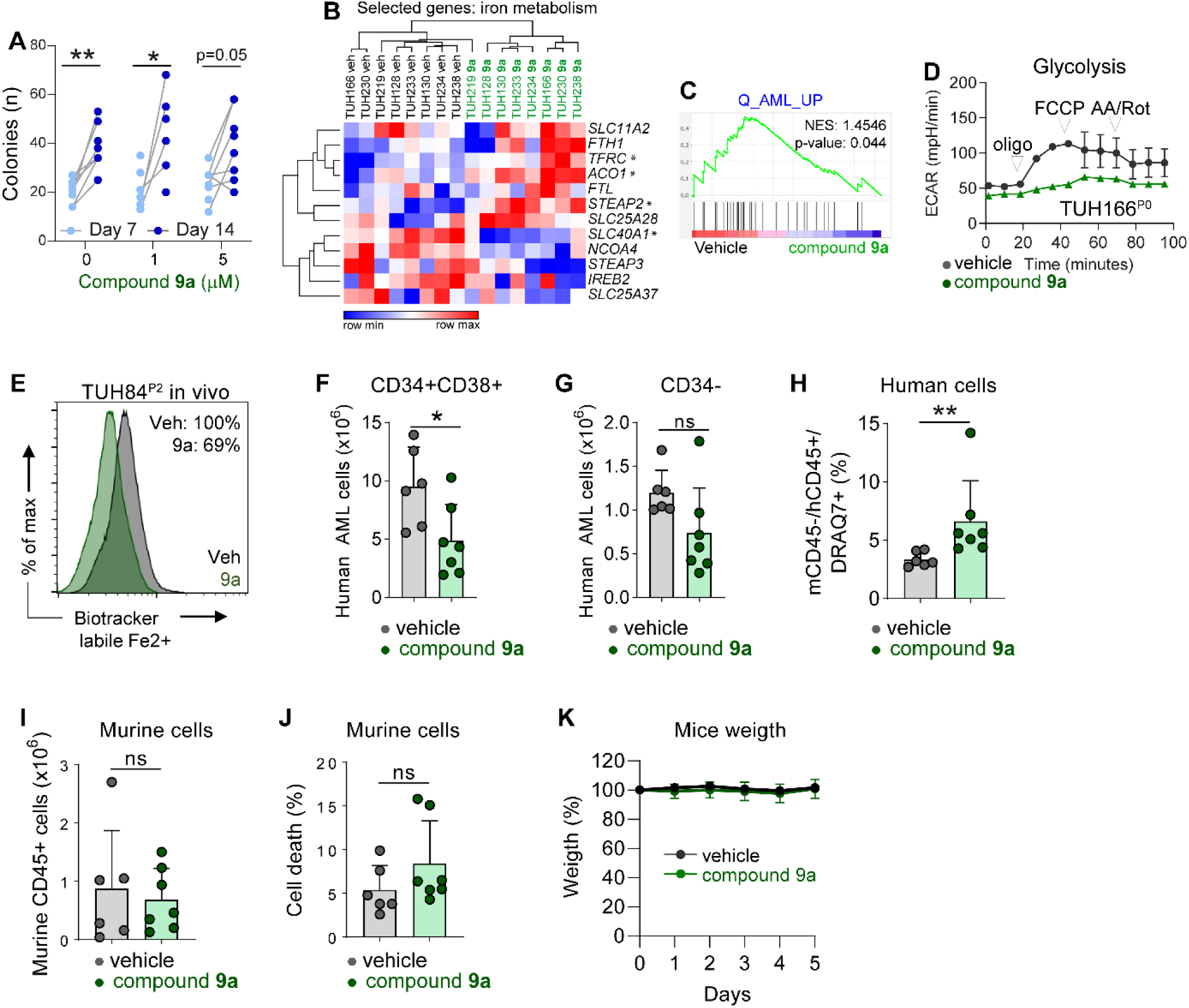
Evaluation of Compound 9a for Anti-Leukemic Activity and Hematological Toxicity. **A.** Normal CD34^+^ hematopoietic progenitor cells were incubated with vehicle, 1 µM or 5 μM compound **9a** in methylcellulose cultures. Colonies were counted under an inverted microscope after 7 and 14 days of culture. **B-C.** Primary AML cells were cultured with vehicle or 1 µM compound **9a** for 24 h and a transcriptomic analysis was carried out (n=8). **B.** Hierarchical clustering of selected genes related to iron metabolism. **C.** Enrichment analysis with the Q_AML_UP signature comparing vehicle or compound **9a** conditions. **D.** Primary AML cells were cultured with vehicle or 1 µM compound **9a** for 48 h. Bioenergetic assays were conducted to measure the extracellular acidification rate (ECAR) in mpH/min.. Oligomycin (Oligo, mitochondrial ATP synthase inhibitor), Carbonyl cyanide-4 (trifluoromethoxy) phenylhydrazone (FCCP, mitochondrial uncoupling agent), and rotenone and antimycin (AA/Rot, complex I and complex III inhibitors, respectively) were used during the assays. **E-K.** Leukemic cells were propagated in NSG mice for 12 weeks. Subsequently, mice received either placebo or 10 mg/kg compound **9a** via daily intraperitoneal injection for five consecutive days. **E.** Intracellular Fe2^+^ quantification using the BioTracker Labile Fe2^+^ dye. **F-G.** Burden of human CD45^+^CD33^+^ leukemic cells within the CD34^+^CD38^+^ (**F**) and CD34^-^ (**G**) compartments. **H.** Measurement of cell death using DRAQ7^+^ staining within hCD45^+^hCD33^+^ human leukemic cells. **I.** Quantification of murine mCD45^+^ hematopoietic cells. **J.** Measurement of cell death using DRAQ7^+^ within murine hematopoietic cells. **K.** Evolution of mice weight (in percent of their initial weight) during treatment with placebo or compound **9a**. Error bars indicate standard deviations. "ns" denotes no significance, *p<0.05, **p<0.01.

## Supplementary Data Files Legends (Excel Files)

**Data File S1. LSC signatures from literature (related to** Figure 2). The list of genes defining the following molecular signatures is provided. LSC104_UP, LSC104_DOWN, LSPC-Quiescent, LSPC-Primed-Top100, LSPC-Cycle-Top100, GMP-like-Top100, ProMono-like-Top100, Mono-like-Top100, cDC-like-Top100.

**Data File S2. Gene sets defining the molecular signatures developed in this study (related to** Figure 2). Q_AML_UP: quiescence signature in AML (bulk); C_AML_UP: cycling signature in AML (bulk); q_cluster genes: genes only regulated in the cluster of cells defined by the Q_AML_UP signature (scRNA-seq); Q_AML_UP ∩ q_cluster: intersection of Q_AML_UP and q_cluster genes; QS35: quiescence signature built from the q_cluster genes in AML cohort data sets.

